# Synapse specific and plasticity-regulated AMPAR mobility tunes synaptic integration

**DOI:** 10.1101/2024.03.19.584837

**Authors:** Agata Nowacka, Angela M. Getz, Hanna L. Zieger, Diogo Bessa-Neto, Christelle Breillat, Sophie Daburon, Cécile Lemoigne, Sébastien Marais, Mathieu Ducros, Andrew C. Penn, Matthieu Sainlos, Daniel Choquet

## Abstract

Synaptic responses adapt to fast repetitive inputs during bursts of neuronal network activity over timescales of milliseconds to seconds, either transiently facilitating or depressing. This high-frequency stimulus-dependent short-term synaptic plasticity (HF-STP) relies on a number of molecular processes that collectively endow synapses with filtering properties for information processing, optimized for the transmission of certain input frequencies and patterns in distinct circuits ^1–3^. Changes in HF-STP are traditionally thought to stem from changes in pre-synaptic transmitter release ^1,2^, but post-synaptic modifications in receptor biophysical properties or surface diffusion also regulate HF-STP ^4–11^. A major challenge in understanding synapse function is to decipher how pre- and post-synaptic mechanisms synergistically tune synaptic transmission efficacy during HF-STP, and to determine how neuronal activity modifies post-synaptic signal computation and integration to diversify neuronal circuit function. Here, taking advantage of new molecular tools to directly visualize glutamate release ^12^ and specifically manipulate the surface diffusion of endogenous AMPAR in intact circuits ^13^, we define the respective contributions of pre-synaptic glutamate release, AMPAR desensitization and surface mobility to frequency-dependent synaptic adaptation. We demonstrate that post-synaptic gain control and signal integration capacity in synaptic networks is influenced by synapse-specific differences in AMPAR desensitization and diffusion-trapping characteristics that are shaped by molecular signaling events recruited during LTP.

## Introduction

The efficacy of synaptic transmission is defined by pre-synaptic transmitter release properties and the number, nanoscale molecular organization, and gating properties of post-synaptic receptors ^14,15^. The prevailing view is that rapid changes in synaptic transmission efficacy during HF-STP are mediated mostly by pre-synaptic mechanisms, through an increase or decrease in the number of transmitter quanta released by successive action potentials ^1,2^. However, post-synaptic mechanisms that rapidly modify the number of receptors available to be activated during repetitive stimulations have also been shown to shape synaptic response amplitudes during HF-STP ^4–11^.

This is particularly important for the frequency-dependent adaptation of fast excitatory transmission mediated by low affinity AMPA-type glutamate receptors (AMPAR), the major excitatory neurotransmitter receptor in the brain. High-density clusters of AMPAR are positioned opposite transmitter release sites, and the probability of receptor activation drops rapidly with their distance from sites of vesicle fusion ^16–19^. Within synapses, diffusing AMPAR are transiently stabilized by the interactions of their auxiliary subunits with scaffolding proteins in the post-synaptic density (PSD) ^7,15,20^. Through the process of Brownian diffusion, AMPAR undergo fast movements, covering tens of nanometers over tens of milliseconds ^4,21,22^. These movements are fast and extensive enough to reorganize receptors positioned in front of glutamate release sites during time intervals relevant for HF-STP. AMPAR mobility has been proposed to contribute to accelerate the recovery of depressing post-synaptic response amplitudes during trains of stimulations, by allowing for the replacement of receptors in the desensitized state (lasting up to a few hundreds of ms) by naïve ones ^4,14^. A similar phenomenon has been observed for Kainate receptors ^23^. This process is thought to be amplified by the faster diffusion of desensitized receptors as compared to naïve ones, as reported at both excitatory ^6^ and inhibitory synapses ^24^ in cultured hippocampal neurons. However, this has not yet been established in more physiological systems such as brain slices due to the lack of adequate tools, and the strict post-synaptic origin of AMPAR immobilization-induced changes in HF-STP has also been questioned ^15^.

The diffusion-trapping equilibrium of synaptic AMPARs is regulated by a variety of stimuli ^4,7,21,25,26^. In particular, calcium influx and consecutive CaMKII activation, as recruited during the induction of Long Term Potentiation (LTP), trigger AMPAR immobilization at synaptic sites ^4,7,26^, and could thus regulate HF-STP as a consequence of synaptic activity history. More generally, control of AMPAR mobility may act as an input-specific gain control mechanism to dynamically adjust the post-synaptic computation of inputs arising from different afferent neurons or during different network states ^8,13,26–28^. Furthermore, given that AMPAR desensitization kinetics ^10,29^ and diffusion-trapping ^7,20,30^ are influenced by their association with a variety of auxiliary subunits that exhibit different expression profiles across the brain, this points to a potentially significant source of post-synaptic variability for the frequency-dependent adaptation of synaptic transmission efficacy ^3,10,11,31–34^. We thus investigated the respective contributions of AMPAR mobility, desensitization and glutamate release to HF-STP at two different synapse types.

## Results

### AMPAR surface diffusion and desensitization regulate HF-STP

HF-STP at excitatory glutamatergic synapses is influenced by both pre- and post-synaptic factors. However, the contribution of post-synaptic AMPAR properties has been relatively understudied (**Fig. 1A**). We first investigated the impact of pre-synaptic glutamate release, AMPAR desensitization, and surface mobility on shaping HF-STP in intact neuronal networks at the canonical central synapse, Schaffer collateral (SC) input to pyramidal neurons (PNs) in the hippocampal CA1 subfield, where the majority of surface AMPARs contain the GluA2 subunit ^31^. To manipulate AMPAR surface mobility *in situ,* we took advantage of the AP-GluA2 knock-in (KI) mouse line we recently developed ^13^, in which the AMPAR GluA2 subunit is tagged with a N-terminal biotin Acceptor Peptide tag (AP-tag). Regulated expression of biotin ligase (BirA^ER^) allows cell-specific biotinylation of AP-GluA2 and expression of biotinylated receptors on the cell surface. A short incubation (20 min) with the tetravalent avidin biotin binding protein NeutrAvidin (NA), which binds and cross-links neighboring biotinylated receptors, allows specific surface immobilization of biotinylated AP-GluA2-containing AMPARs (bAP-GluA2). Alternatively, incubation with a monovalent avidin, monomeric StreptAvidin (mSA), allows live tracking of surface receptor mobility ^13,35^ (**Fig. 1B**). BirA^ER^-eGFP was introduced into CA1 PNs of organotypic hippocampal slices (OHS) via single cell electroporation (SCE). Lattice light sheet microscopy (LLSM) imaging confirmed efficient tissue penetration and specific binding of fluorescent NA to bAP-GluA2 on CA1 PNs expressing BirA^ER^ (**Fig. 1C**), consistent with our previous results ^27^.

**Figure 1.**
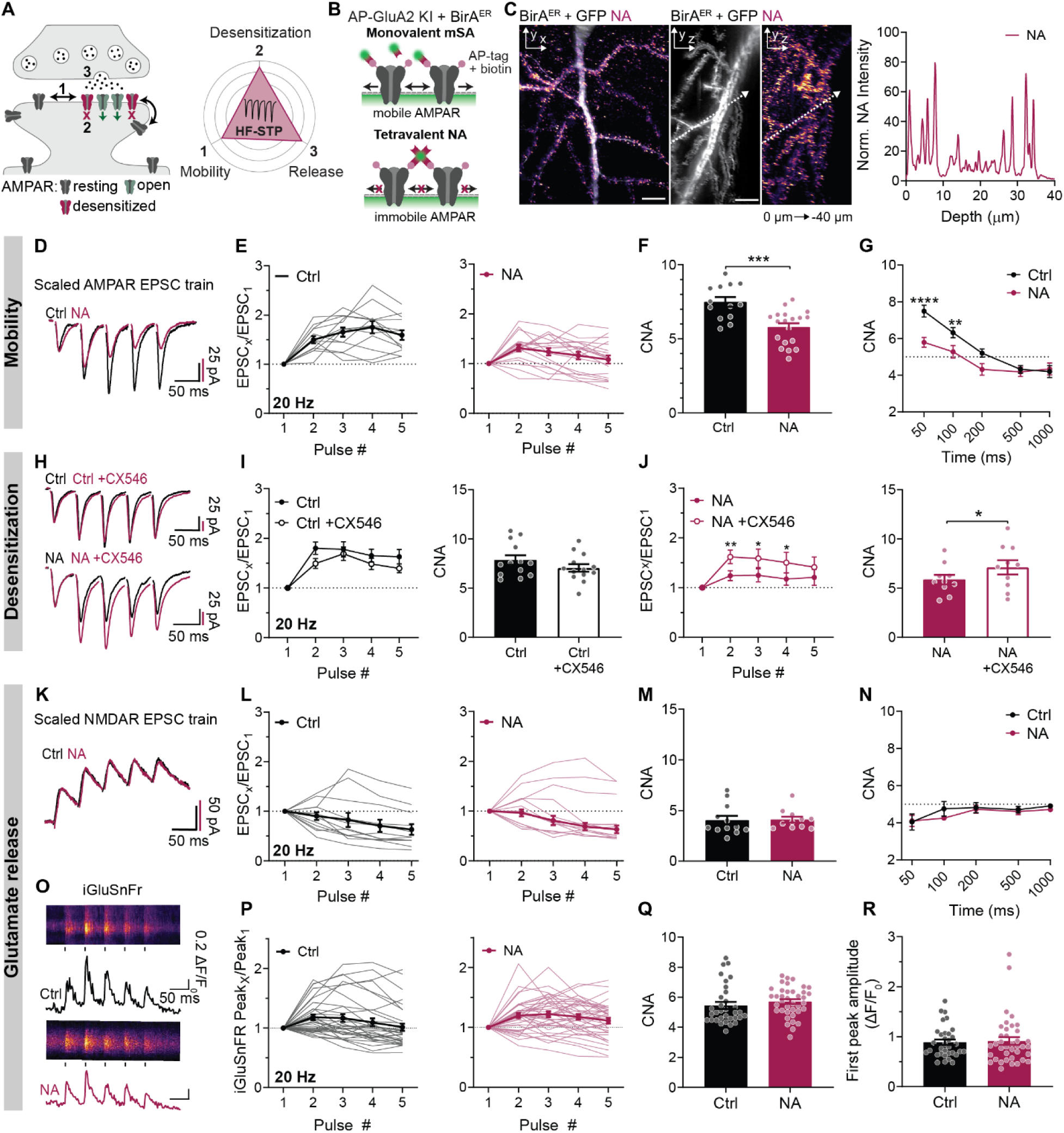
AMPAR surface diffusion and desensitization regulate HF-STP. (**A**) Schematic of pre- and post-synaptic contributions to HF-STP. (**B**) AP-GluA2 KI molecular toolkit to track and manipulate AMPAR surface mobility. (**C**) LLSM images of bAP-GluA2 labelled with NA, CA1 PN in OHS, and NA intensity profile in depth. Scale bar, 10 μm. (**D**) Exemplary AMPAR EPSC trains (20 Hz) in SC – CA1 ± NA cross-link. (**E**) Normalised AMPAR EPSC amplitudes ± NA (n= 13, 19 cells). Data on all graphs represented as scaled or normalised to first-peak amplitude and Mean ± SEM in bold unless otherwise stated. (**F**) Data from **E** as cumulative normalised amplitude (CNA) (***P=0.0003; Unpaired t-test). (**G**) CNA of trains ± NA with varying ISI (n=8-15 and 9-19 cells; F=13.48, P= 0.0004, **-****P≤0.0074; Two-way ANOVA, Fisher’s LSD post-hoc test). (**H**) Exemplary AMPAR EPSC trains ± NA and ± CX546. (**I** - **J**) Normalised AMPAR EPSC amplitudes as in **H** (n=13, 10 cells; F=2.367, P=0.1499; F=10.22, P=0.0109, *-**P≤0.0313; Two-way ANOVA, Fisher’s LSD post-hoc test (left); P=0.1499, *P=0.0109; Paired t-test (right)). (**K**) Exemplary NMDAR EPSC trains ± NA. (**L** - **M**) Normalised NMDAR EPSC amplitudes and corresponding CNA (n= 12, 11 cells; P=0.4134; Mann-Whitney test). (**N**) CNA of NMDAR EPSC trains with varying ISI (n=11-12 and 10-13 cells; P≥0.3867; Mann-Whitney test). (**O**) Representative SF-iGluSnFR-A184V transients in SC-CA1 dendritic spines during 20 Hz trains ± NA. (**P**) Average normalized iGluSnFR response per spine, (n= 33, 38 spines; 3-10 trials). (**Q**) Data from **P** represented as CNA (P=0.0869; Mann-Whitney test). (**R**) iGluSnFR first-peak amplitudes ± NA (P=0.8143; Mann-Whitney test).

Having confirmed successful application of our molecular toolkit to control and image AMPAR mobility in brain slices, we performed whole-cell voltage-clamp recordings of CA1 PNs following NA or vehicle incubation. Trains of 5 electrical stimulations at 20 Hz were applied to the SC to induce HF-STP. The recordings were performed in elevated extracellular Ca^2+^ (4 mM) to enhance pre-synaptic glutamate release probability (P_r_) and thereby minimize the pre-synaptic variable. We measured the amplitudes of evoked AMPAR EPSCs (**Fig. 1D - E**) induced by the stimulation trains, and observed enhanced synaptic depression at the SC – CA1 synapse in OHS following NA cross-link (**Fig. 1F**) (22.7% decrease in cumulative normalized amplitude [CNA] of the trains after NA cross-link as compared to control conditions). This enhancement was observed at 50 and 100 ms interstimulus interval (ISI), but was absent with ISI ≥ 200 ms (**Fig. 1G**). A similar, though less pronounced, enhancement in synaptic depression was observed at lower extracellular Ca^2+^ concentration (2.5 mM) (**Supplementary Fig. 1**). This is to be expected, as in lower extracellular Ca^2+^, the P_r_ is decreased, thus the time between two sequential vesicle release events at the same site is increased, allowing for receptors to recover from desensitization between release events and reducing the impact of surface diffusion ^14^. Additionally, consistent with previous findings ^13,27^, AMPAR cross-link had no effect on basal synaptic transmission (**Supplementary Fig. 2**).

Depression at the SC – CA1 synapse of OHS following NA cross-link was prevented by the application of the AMPAR desensitization blocker CX546 (**Fig. 1F - J**). This indicates that the enhanced synaptic depression induced by receptor immobilization is dependent on AMPAR desensitization. It is consistent with the diffusion-dependent exchange of AMPAR that normally acts to reduce the number of desensitized AMPARs positioned in front of glutamate release sites. To avoid potential pre-synaptic effects of chemical desensitization blockers, we examined the impact of NA cross-link on HF-STP in CA1 PNs expressing AMPAR desensitization mutants. The stabilization of the closed clamshell conformation of the receptor contributes to the stability of the ET/YR (E713T, Y768R) AMPAR mutant in the desensitized state, resulting in a significant deceleration of the recovery from desensitization ^36^. On the contrary, the stabilization of the dimer interface in the L483Y AMPAR mutants renders the receptor resistant to desensitization ^37^. In wild-type rat OHS, we overexpressed AP-SEP-GluA2 ET/YR or LY along with AP-tdTom-GluA1 and BirA^ER^ (**Supplementary Fig. 3A**). Expression of the ET/YR mutant receptors resulted in a profound increase in synaptic depression after the receptors were cross-linked with NA, visible with ISI up to 1 sec (**Supplementary Fig. 3B - C**). Conversely, in neurons expressing the desensitization resistant L483Y mutant receptors, we did not observe any effect of AMPAR immobilization on HF-STP (**Supplementary Fig. 3D - E**). This again suggests that blocking AMPAR surface diffusion traps desensitized receptors in the synapse and increases synaptic depression during HF-STP. To establish whether NA binding could directly affect AMPAR gating, we performed fast glutamate application onto outside-out patches pulled from the cell membranes of HEK293T cells expressing AP-GluA2-BirA^ER^ and GluA1, or nucleated outside-out patches form CA1 PNs in OHS expressing BirA^ER^-eGFP or eGFP alone. NA binding to bAP-GluA2 had no effect on AMPAR desensitization kinetics in both models (**Supplementary Fig. 4**).

AMPAR cross-link with NA could potentially affect synaptic nano-organization ^16,17^, thereby impacting the fidelity of high-frequency synaptic transmission. To assess changes in AMPAR nano-organization, we performed dSTORM imaging in dissociated rat hippocampal neuronal cultures overexpressing AP-mRFP-GluA1, AP-SEP-GluA2 and BirA^ER^. We found no change in the number of AMPARs per spine, AMPAR cluster number or AMPAR numbers per cluster after NA binding (**Supplementary Fig. 5A – E**). Notably, with uPAINT imaging, we observed a strong decrease in the mobile fraction of AMPARs upon application of NA, confirming its immobilizing effect on the dynamic organization of AMPARs (**Supplementary Fig. 5F – H**).

Traditionally, HF-STP regulation is attributed primarily to pre-synaptic mechanisms ^1,2^. To confirm that NA affects HF-STP solely through post-synaptic retention of desensitized AMPARs at synapses, we examined whether AMPAR cross-link could impair synaptic facilitation by indirectly modulating pre-synaptic glutamate release. NA application showed no effect on isolated NMDAR current trains evoked by 20 Hz stimulations, indicating no change in pre-synaptic glutamate release (**Fig. 1K – N)**. To further probe for potential changes in glutamate release in response to AMPAR cross-link, we directly measured glutamate release events during HF-STP trains using fast 2-photon linescan imaging with SF-iGluSnFr-A184V, a genetically encoded glutamate sensor ^12^. To resolve individual spines, we transduced CA1 neurons in OHS from AP-GluA2 KI mice with a combination of AAVs encoding BirA^ER^ and Cre recombinase with FLEx SF-iGluSnFR-A184V, to obtain sparse iGluSnFR expression only in neurons expressing bAP-GluA2 (**Supplementary Fig. 6**). Sequential linescans through activated dendritic spines were performed to measure iGluSnFR fluorescence transients during 20 Hz stimulations evoked by an extracellular stimulating electrode. NA cross-link of surface AMPARs had no effect on glutamate release characteristics during HF-STP trains (**Fig. 1O - Q**), and there was no change in the first-peak amplitude of iGluSnFR transients, suggesting no alteration in pre-synaptic release probability (**Fig. 1R**). Altogether, this indicates that the enhanced synaptic depression following AMPAR cross-link is expressed through post-synaptic mechanisms and is not associated with decreased pre-synaptic glutamate release during successive stimuli.

In summary, these data demonstrate that AMPAR surface diffusion sustains synaptic transmission amplitudes during high-frequency activity, by allowing replacement of desensitized receptors by naïve ones in front of glutamate release sites.

### Differential AMPAR biophysics and mobility contribute to synapse-specific HF-STP

HF-STP phenotypes vary between brain regions, synapse types and even synapses on the same neuron (e.g. ^33,38^, reviewed in ^1–3,39^). This diversity relies on both pre- and post-synaptic mechanisms such as differences in release probability as well as the molecular organization of the post-synapse, including AMPAR dynamics ^4^. Our initial data were obtained at the prototypical SC – CA1 synapses that display a low and heterogenous P_r_ ^40,41^ along with a modest facilitation during HF-STP. We reasoned that the impact of AMPAR mobility on HF-STP would be more pronounced at synapses with high Pr, where the extent of receptor recovery from desensitization between sequential release events at the same location would be reduced. We therefore compared the effect of AMPAR cross-link on HF-STP and pre-synaptic glutamate release at the SC - CA1, and L4 - L2/3 synapses of the barrel field of the primary somatosensory cortex (S1). The latter displays a high P_r_ ^42^ and strong synaptic depression, which contributes to adaptation of the synapse to incoming sensory inputs ^43^. Moreover, at L2/3 synapses, AMPAR surface trafficking is known to play an important role in regulation of synaptic plasticity ^28,44^. To analyze the role of AMPAR immobilization on HF-STP in experimental models closer to physiological conditions, we performed the experiments in *ex vivo* brain slices from adult AP-GluA2 KI mice stereotaxically injected with AAVs encoding BirA^ER^-Cre and FLEx eGFP into selected brain regions (**Fig. 2A - B**). The recordings were performed in normal extracellular Ca^2+^ conditions (2.5 mM). We confirmed NA penetration into acute brain slices and specific binding to biotinylated receptors with LLSM (**Fig. 2C**).

**Figure 2.**
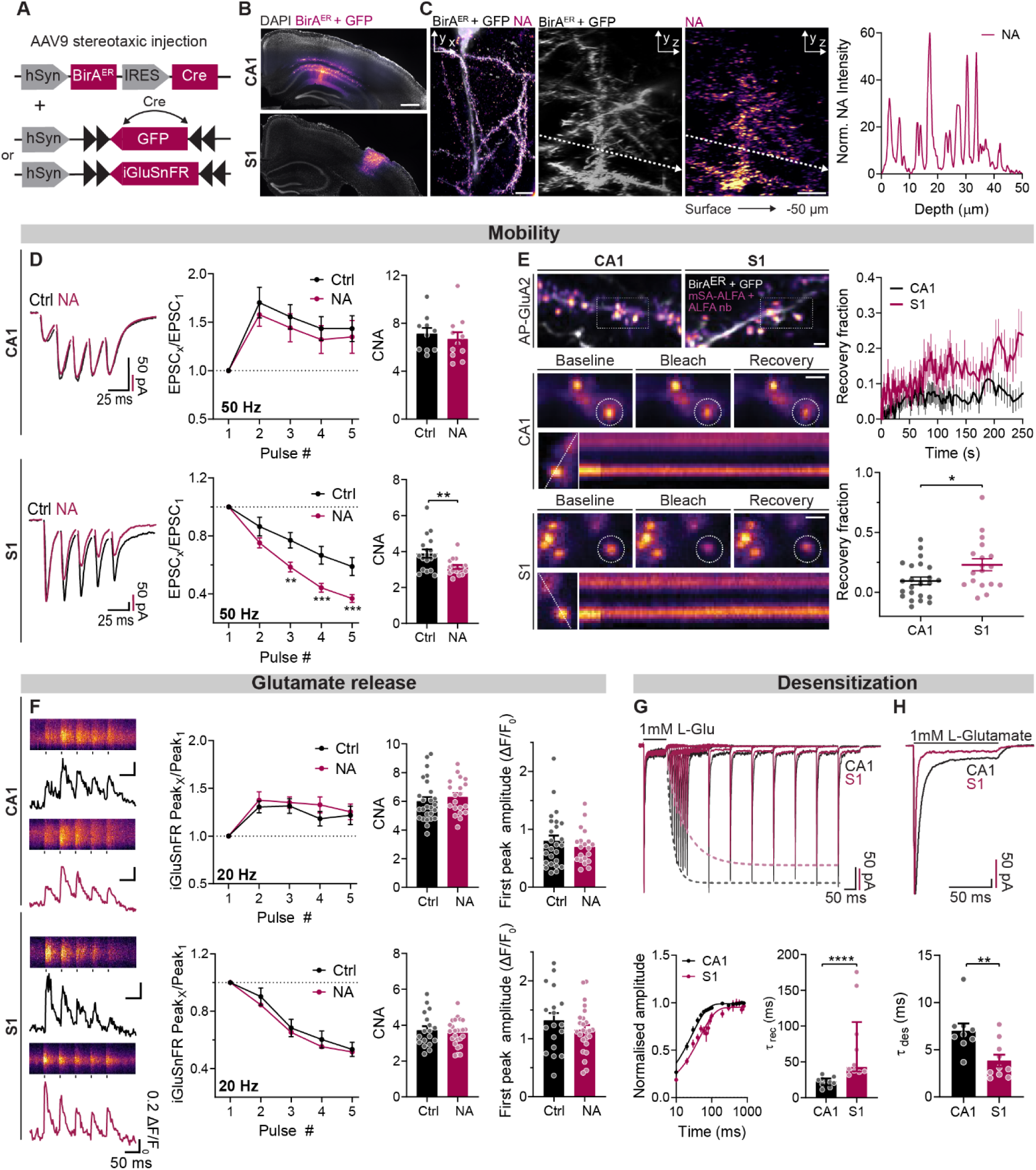
Differential AMPAR biophysics and mobility contribute to synapse-specific HF-STP. (**A**) AAV9 constructs for stereotaxic injections. (**B**) Representative injections in CA1 and S1. Scale bar, 500 μm. (**C**) LLSM images of bAP-GluA2 labelled with NA, CA1 PN in acute slice, and NA intensity profile in depth. Scale bar, 10 μm. (**D**) Exemplary AMPAR EPSC trains (50 Hz) in SC – CA1 and S1 L4 – L2/3 ± NA (n= 10, 11 and 18, 17 cells; F=0.3460, P=0.5633; F=8.243, P=0.0071, **-***P≤0.0034; Two-way ANOVA, Fisher’s LSD post-hoc test) and corresponding CNA (P=0.5633, Unpaired t-test; **P=0.007, Welch’s test). (**E**) LLSM-FRAP in acute slices, bAP-GluA2 labelled with mSA-ALFA and αALFA nanobody (Atto 643). Representative images show spine ROIs (dashed circles) at baseline (−1 s), after photobleaching (+0.5 s) and diffusion-dependent recovery (+250 s). Scale bar, 2 μm. Kymographs illustrate ROI fluorescence recovery profiles (dashed line; ∼250 s). Mean recovery curves and fraction in CA1 and S1 (n= 23, 18 spines; *P=0.0240; Unpaired t-test). (**F**) Representative SF-iGluSnFR-A184V transients in CA1 and S1 dendritic spines during 20 Hz trains ± NA. Average normalized iGluSnFR response per spine (n= 26, 20 and 18, 25 spines; 3-10 trials; F=0.55127, P=0.4777; F=0.3779, P=0.5421; Two-way ANOVA, Fisher’s LSD post-hoc test), and corresponding CNA (P=0.3060, Mann-Whitney test; P=0.5421, Unpaired t-test) and first-peak amplitudes (P=0.6847, Mann-Whitney test; P=0.2768; Unpaired t-test). (**G**) Exemplary AMPAR recovery from desensitization curves during glutamate applications with increasing ISI, somatic nucleated outside-out patches from CA1 or S1 PNs (n=10 cells). AMPAR rate of recovery from desensitization (Median ± quartiles; ****P<0.0001, Mann-Whitney test). (**H**) AMPAR rate of desensitization (n= 9, 10 cells; **P=0.0054, Unpaired t-test).

At the SC–CA1 synapse in *ex vivo* brain slices, AMPAR cross-link did not induce a detectable effect on HF-STP, regardless of pulse train interstimulus intervals (**Fig. 2D**) (**Supplementary Fig. 7A**). A modest decrease in synaptic facilitation upon NA-induced cross-link was observed in increased extracellular Ca^2+^ concentration (4 mM), as expected from a lower inter-release interval at given synapses in these conditions (**Supplementary Fig. 8**). On the contrary, AMPAR cross-link in L4-L2/3 synapses in S1 profoundly enhanced synaptic depression in regular extracellular Ca^2+^ conditions (2.5 mM) (**Fig. 2D**) (**Supplementary Fig. 7B**). We reasoned that the synapse-specific difference in AMPAR immobilization on HF-STP could derive from intrinsic differences between the two cell types in AMPAR surface diffusion rates or entry/exit rates from the desensitized states. To measure the diffusion of surface AMPARs in acute slice preparations, we expressed the Cre-dependent BirA^ER^ in both brain regions and used LLSM, combined with FRAP, of AMPARs labelled with mSA-ALFA and Atto 643-conjugated ALFA-nanobody ^45^ (**Fig. 2E**). We found that the synaptic AMPAR recovery fraction was ∼ 2.5-fold higher at spines on basal dendrites of L2/3 PNs (receiving vertical inputs from L4) in S1 (23.35%) than at spines on apical dendrites in the *stratum radiatum* of CA1 PNs (9.55%) (**Fig. 2E**), indicating distinct, synapse-specific diffusion-trapping equilibriums for synaptic AMPARs. The higher diffusion rate of AMPARs in S1 is consistent with the stronger impact of AMPAR cross-link on HF-STP at these synapses.

In parallel, we used fast 2-photon linescan imaging with SF-iGluSnFr-A184V at dendritic spines in S1 and CA1 (as above) to resolve potential pre-synaptic contributions to HF-STP. In control conditions, we observed facilitation of iGluSnFR fluorescence transients during HF-STP trains (20 Hz) in SC – CA1, and depression in L4 – L2/3, demonstrating that glutamate release characteristics are different between the two synapse types, and consistent with the corresponding electrophysiological HF-STP profiles. NA cross-link did not impact pre-synaptic glutamate release profiles measured with iGluSnFR (**Fig. 2F**). In addition, the first-peak amplitude of iGluSnFR transients was higher at S1 than CA1 synapses, in agreement with the higher P_r_, and was not affected by NA cross-link at either synapse type. Altogether, these data demonstrate that synapse-specific AMPAR mobility participates to gain control during HF-STP through a post-synaptic mechanism.

The other post-synaptic components which may account for the differential impact of AMPAR cross-link on HF-STP in CA1 and S1 are the desensitization kinetics of AMPAR complexes. Indeed, these exhibit variability across cell types and depend on AMPAR subunit and auxiliary protein composition ^10^. Fast glutamate application on somatic nucleated outside-out patches pulled from CA1 and S1 pyramidal neurons revealed significantly slower AMPAR recovery from desensitization in S1 compared to CA1 PNs (τ_rec_ 71.45 vs 22.3 ms) (**Fig. 2G**). Moreover, AMPARs on PNs from the S1 desensitized approximately 2-fold faster than those from the CA1 (τ_des_ 3.854 vs 7.012 ms) (**Fig. 2H**).

Collectively, these findings demonstrate that the unique characteristics of HF-STP at each synapse are determined by a blend of inherent differences in pre-synaptic release probability, AMPAR mobility, and desensitization kinetics.

### CaMKII activity and LTP induction modulate AMPAR diffusion-trapping and tune HF-STP

Following LTP-like induction paradigms, extrasynaptic AMPARs are recruited to synaptic sites through diffusion-trapping mechanisms, involving CaMKII-dependent phosphorylation of AMPAR auxiliary proteins for their increased stabilization at synapses ^4,7,21,26,30^. We thus investigated if this physiological mechanism of AMPAR immobilization by diffusion-trapping could tune HF-STP during LTP. We induced LTP in the L4–L2/3 pathway in S1 of *ex vivo* brain slices from AP-GluA2 KI mice with a protocol pairing post-synaptic depolarization (0 mV) and low frequency stimulation (0.167 Hz for 5 mins). Stable LTP (147%) was achieved (**Fig. 3A - B)**, and coincided with a transient enhancement of synaptic depression during HF-STP trains (50 Hz) 5 and 10 minutes after the onset of LTP induction, with no change in control conditions (no pairing, stimulation rate kept at 0.033 Hz) (**Fig. 3 B – D)**. HF-STP profiles returned to baseline at later timepoints. This is consistent with the models of AMPAR trafficking during LTP, where activity-dependent trapping of AMPARs is critical during the early phases of LTP (eLTP), whereas in the later phases, highly mobile exocytosed AMPARs allow potentiated synapses to recover a normal fraction of mobile receptors ^30^.

**Figure 3.**
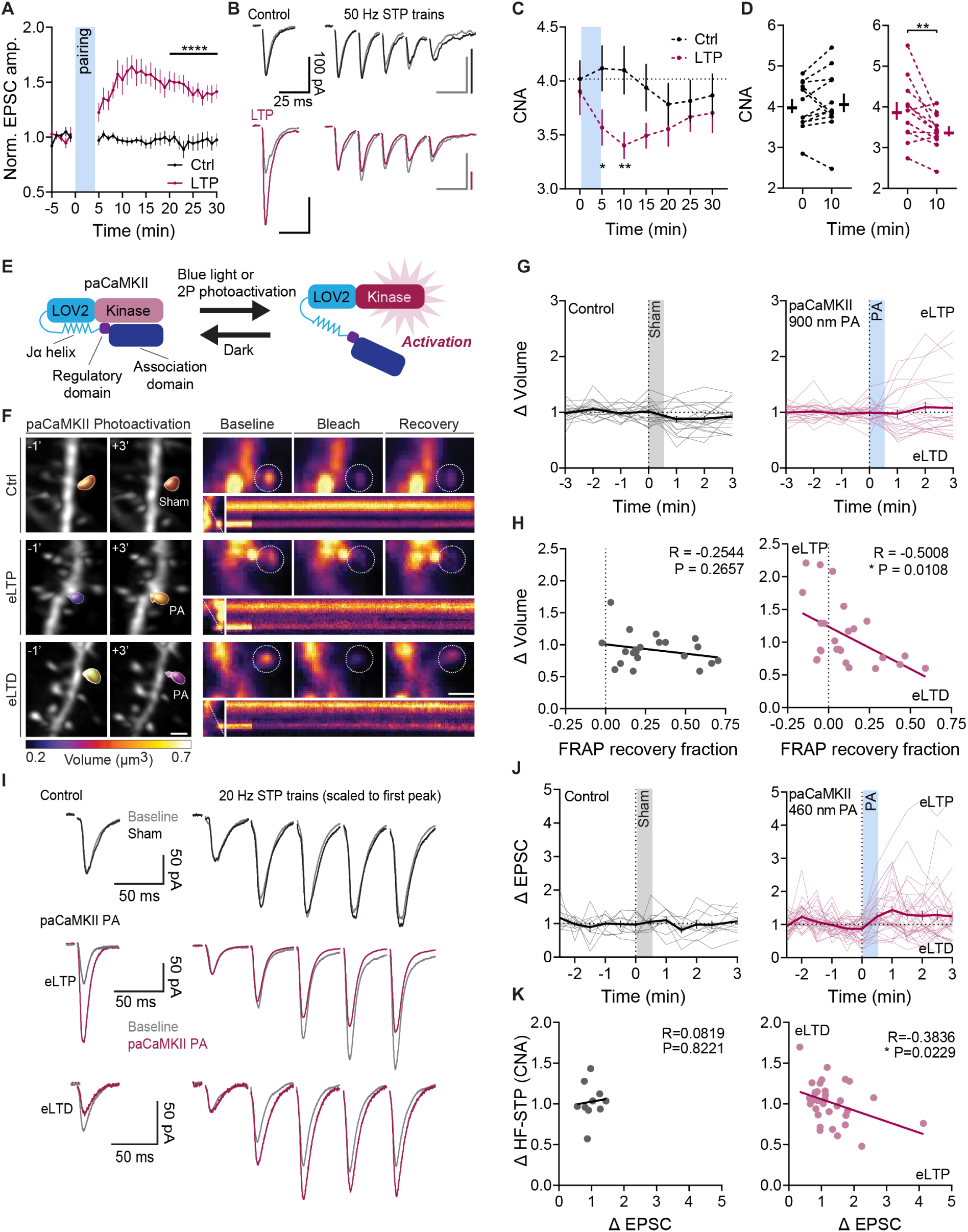
CaMKII activity and LTP induction modulate AMPAR diffusion-trapping and tune HF-STP. (**A**) Normalized AMPAR EPSC amplitudes from S1 L4 - L2/3 PNs during electrical LTP induction in acute slices (n=12 cells; ****P<0.0001, Unpaired t-test). (**B**) HF-STP trains (50 Hz) were acquired every 5 min during LTP recordings. Representative evoked AMPAR EPSC and trains before (0 min) and after LTP induction (10 min). (**C**) HF-STP profiles ± LTP induction (F=2.332, P=0.0358, *-**P≤0.0375; Two-way ANOVA, Fisher’s LSD post-hoc test). (**D**) Change in CNA between baseline (0 min) and 10 min (P=0.5243, **P=0.0087, Paired t-test). Plotted individual paired values and mean ± SEM (**E**) paCaMKII construct based on the LOV2 photosensitive domain. Adapted from ^47^. (**F**) Representative 3D reconstructions of spine volume changes during eLTP/eLTD and corresponding FRAP measurements of AMPAR surface diffusion 3 min after photoactivation (circles; baseline -1 s, bleach + 1 s, recovery + 240 s). Kymographs illustrate FRAP rate across the acquisition (dashed lines; ∼250 s). (**G**) Normalized spine volume changes over time (n= 21, 25 spines). (**H**) Cross-correlation of paCaMKII-induced structural plasticity and AMPAR FRAP recovery fraction. eLTP enhances, whereas eLTD decreases AMPAR synaptic trapping. (**I**) Exemplary AMPAR EPSC amplitudes and HF-STP trains (20 Hz) at SC - CA1 in OHS before (0 min) and after paCaMKII photoactivation (3 min). (**J**) Normalised AMPAR EPSC amplitudes during recordings (n= 10, 35 cells). (**K**) Cross-correlation of paCaMKII-induced plasticity of synaptic transmission efficacy (eLTD/eLTP) and HF-STP CNA. eLTP enhances, whereas eLTD decreases short-term depression.

At this stage, there remained the possibility that the changes in HF-STP during the first minutes following LTP induction could result from pre-synaptic phenomenon ^46^. To further investigate the role of AMPAR diffusion-trapping in tuning HF-STP during eLTP, we took advantage of photoactivatable CaMKII (paCaMKII) - an optogenetic form of CaMKIIα fused to the LOV2 photosensitive domain, activated by blue light or 2-photon excitation ^47^ - that we expressed along with BirA^ER^ in CA1 PNs of OHS from AP-GluA2 KI mice (**Fig. 3E**). Photoactivation of paCaMKII ensures a purely post-synaptic origin of synaptic potentiation, which, like electrically-induced LTP, is associated with spine volume changes that can be used as a readout of long-term plasticity ^47^. We followed spine volume changes for 3 min following photo-activation (PA), corresponding to the eLTP phase. Without PA, we observed no change in spine volume, whereas with a train of spine-targeted 2-photon laser pulses (900 nm, 3 mW, 30 x 80 ms at 1 Hz), we could induce structural plasticity. PA of paCaMKII induced bidirectional plasticity, with increases (eLTP) or decreases (eLTD) in spine volume (**Fig. 3F – G**), consistent with recent reports demonstrating that CaMKII can induce both LTP and LTD depending on the frequency of input signal through differential autophosphorylation levels of the holoenzyme ^48^. We tested whether paCaMKII activation modulates AMPAR diffusion-trapping at Schaffer collateral synapses on *stratum radiatum* dendrites of CA1 PNs, using LLSM-FRAP and labelling of endogenous bAP-GluA2 with mSA-ALFA and Atto 643-conjugated ALFA-nanobody. We found a significant negative correlation between spine volume change and FRAP recovery fraction following photo-activation (**Fig. 3F, H**). Spines that potentiated exhibited reduced AMPAR mobility during eLTP, indicating that CaMKII enhances AMPAR stabilization at synaptic sites through diffusion-trapping, whereas spines that depressed exhibited increased mobility during eLTD. Interestingly, these opposing effects of eLTP and eLTD on AMPAR surface mobility are consistent with previous reports ^4,8^.

We then examined how the regulation of AMPAR diffusion-trapping by paCaMKII affects HF-STP using whole-cell voltage-clamp recordings of evoked AMPAR currents in CA1 PNs expressing paCaMKII and BirA^ER^, using a whole-cell blue light photo-activation paradigm (460 nm, 3 mW, 30 x 80 ms at 1 Hz). We again observed the induction of bidirectional plasticity (eLTP and eLTD), with a significant negative correlation between the directionality of potentiation and changes in HF-STP train profiles (20 Hz). Synapses exhibiting eLTP or eLTD respectively displayed enhanced or reduced depression of synaptic transmission during HF-STP trains (**Fig. 3I-K**), consistent with the effects of paCaMKII activation on enhancing or decreasing AMPAR immobilization. Altogether, these data indicate that the physiological activity patterns that induce synaptic plasticity (LTP/LTD) and recruit CaMKII signaling act to tune AMPAR diffusion-trapping, consequently shaping the dynamic range of post-synaptic responses during high-frequency activity.

### AMPAR immobilization tunes post-synaptic computation and synaptic network integration

We next aimed to define how AMPAR immobilization, and the associated changes in HF-STP, could contribute to signal processing and integration in synaptic networks. HF-STP operates as a mechanism to adapt synaptic gain across different input frequencies during on-going synaptic activity ^49^. In this context, the fine-tuning of post-synaptic Ca^2+^ influx into dendritic spines is critical for maintaining and adapting synaptic transmission, information transfer and storage in neuronal networks. To test whether AMPAR surface mobility regulates spine Ca^2+^ transients during HF-STP, we transduced CA1 PNs in OHS from AP-GluA2 KI mice with BirA^ER^-Cre and FLEx jGCaMP8m and applied NA mediated AMPAR cross-link as a surrogate to activity-induced immobilization. We used 2-photon area scans to measure GCaMP fluorescence transients in individual spines on apical dendrites of CA1 PNs evoked by electrical stimulation of SC (10 x 20 Hz) in regular P_r_ conditions (2.5 mM Ca^2+^). AMPAR immobilization reduced the amplitude of post-synaptic Ca^2+^ transients (**Fig. 4A - D**). This indicates that the regulation of AMPAR mobility at synapses acts as a post-synaptic gain control filter that can dynamically adjust neuronal information processing.

**Figure 4.**
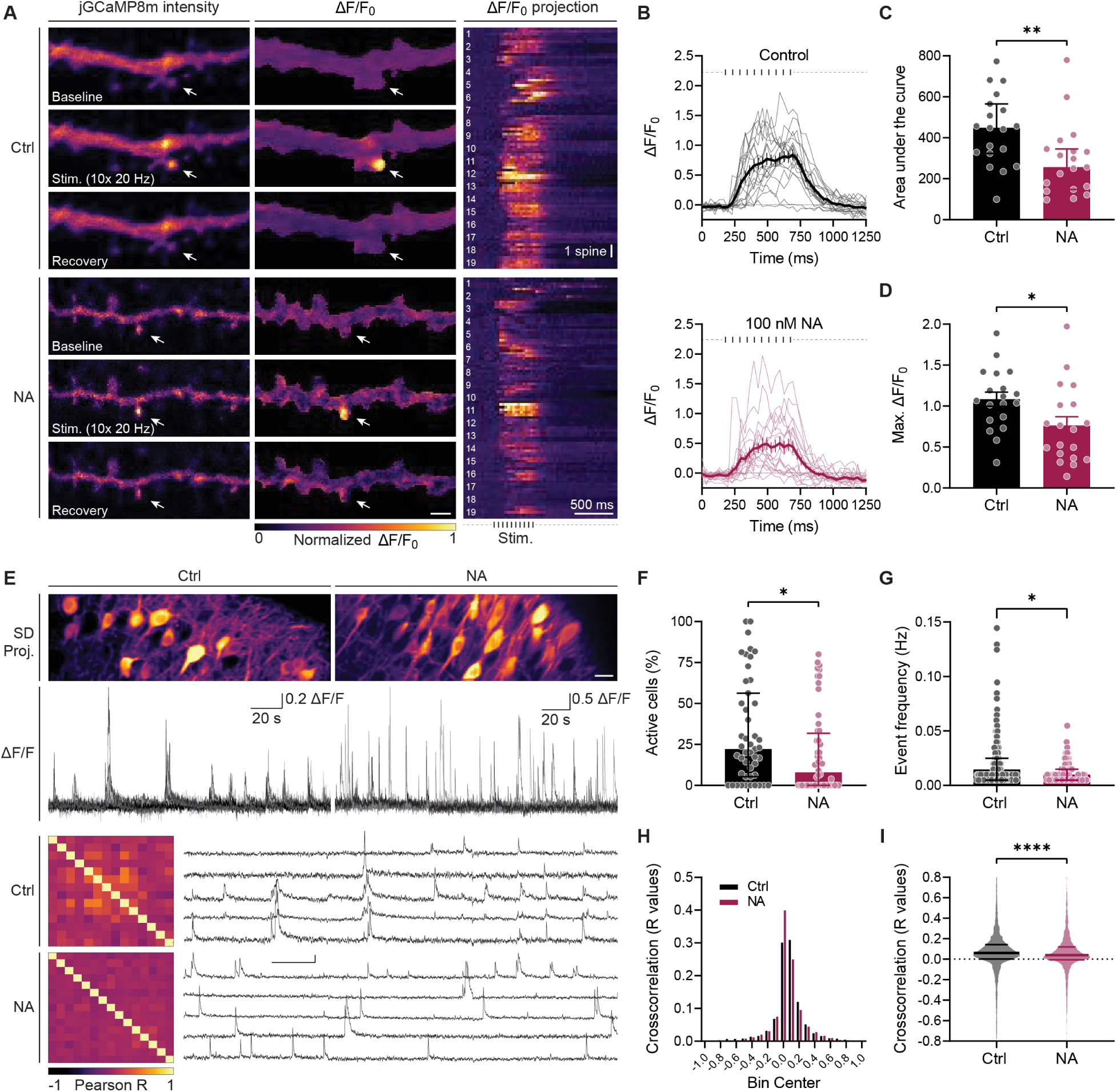
AMPAR immobilization tunes post-synaptic computation and synaptic network integration. (**A**) Representative jGCaMP8m synaptic Ca^2+^ transients in *stratum radiatum* dendrites of CA1 PN in OHS from AP-GluA2 KI during HF-STP trains (20 Hz) ± NA. GCaMP intensity and ΔF/F_0_ averaged projection (left). Arrows indicate active spines. Scale bar, 1 μm. One-pixel averaged projections of spine Ca^2+^ transients (right), each row represents one acquisition and spine replicates are grouped (n= 19 spines, 4-5 trials). (**B**) Spine Ca^2+^ transients during train stimulations ± NA. (**C**, **D**) Corresponding area under the curve (Median ± interquartile; **P=0.0063; Mann-Whitney test) and maximum ΔF/F_0_ (*P=0.0263; Unpaired t-test) of traces in **B**. (**E**) Representative standard deviation projection of jGCaMP8m intensity, FOV of CA1 subfield in OHS from AP-GluA2 KI, and overlay of spontaneous Ca^2+^ transients measured in CA1 PN somas represented as ΔF/F. Representative pairwise cross-correlation matrix of ΔF/F traces and expanded ΔF/F traces of active neurons exhibiting spontaneous events, as above. (**F**) Percentage of active cells per FOV (Median ± quartiles; N=629, 613 cell ROIs, from 52, 47 FOVs in 12, 11 slices; *P=0.0371, Mann-Whitney test) and (**G**) event frequency in active cells ± NA (Median ± quartiles; n= 182, 116 cell ROIs; **P=0.0053, Mann-Whitney test). (**H**, **I**) Histogram and median distribution of Pearson R values from pairwise cross-correlations for cell ROIs, by acquisition FOV (n= 4-29 cell ROIs per FOV; 11 ± 5.67, 12 ± 5.67 (Median ± SD)).

To then examine the contribution of AMPAR mobility to synaptic integration during ongoing network activity, we performed jGCaMP8m area scan imaging to read-out spontaneous cell spiking events within the hippocampal CA1 subfield (**Fig. 4E**). Receptor cross-link decreased the percentage of spontaneously spiking cells (from 12.5% to 3.85%), as well as the spike frequency within the population of active cells (**Fig. 4F - G)**. Moreover, AMPAR cross-link disrupted network synchronicity. In control conditions cells tended to spike in synchronous ensembles, whereas after NA cross-link this synchronicity was lost, as demonstrated by cross-correlation analysis of somatic ΔF/F transients (**Fig. 4H - I**). Altogether, these results demonstrate that AMPAR mobility is critical to sustain and fine-tune synaptic Ca^2+^ integration and signal propagation during high-frequency synaptic transmission, which underlies the activity state and synchronicity of neural networks.

## Discussion

By combining electrophysiology and *in situ* imaging of fluorescent biosensors with recently developed molecular tools to visualize and manipulate AMPAR mobility ^13^, we demonstrate that HF-STP is fine-tuned in a synapse-specific manner by the interdependent contributions of pre-synaptic glutamate release, post-synaptic AMPAR surface diffusion and desensitization kinetics (**Supplementary Fig. 9**). We show that the physiological regulation of AMPAR diffusion-trapping following CaMKII activation and LTP induction modifies HF-STP responses in intact tissue, and we identify roles for AMPAR mobility in tuning Ca^2+^ influx and signal integration during high-frequency synaptic transmission and for regulating neural network activity.

Our results build on a model in which the surface exchange of AMPARs in front of glutamate release sites regulates the extent to which sequential post-synaptic responses are attenuated by the cumulative desensitization of receptors during repetitive synaptic activity. This model predicts that synapses featuring a high probability of neurotransmitter release, along with a significant fraction of AMPARs that are mobile and recover slowly from desensitization, will exhibit the greatest influence of AMPAR mobility on short-term synaptic plasticity. A synapse type that displayed a pronounced contribution of AMPAR diffusion to limiting the impact of AMPAR desensitization on short-term synaptic depression was the layer 4 to layer 2/3 synapses in the primary somatosensory cortex (**Fig. 2D-E**). Synaptic depression is a common feature of transmission at many cortical synapses, and the contribution of AMPAR diffusion may well extend to excitatory synapses onto pyramidal neurons in other areas of the cortex ^50^. Earlier studies did not observe any effect of inhibiting AMPAR desensitization on short-term depression at the layer 4 to layer 2/3 synapses in primary visual cortex ^50^. This is despite the fact that AMPARs desensitize significantly to brief glutamate exposure and recover to the resting state very slowly in excised membrane patches from the same cell-type ^51^, a finding which we also reproduced in our own data (**Fig. 2H**). We propose that the rapid exchange of desensitized synaptic AMPAR by surface diffusion offers an explanation to reconcile these observations, and is consistent with AMPAR immobilization trapping desensitized AMPARs at synapses, thereby enhancing short-term synaptic depression (**Fig. 1D-E**) ^4^. This is also consistent with our observation that mutations in AMPARs that increase the extent and duration to which synaptic AMPARs desensitize only differ in their HF-STP when AMPARs are immobilized (**Supplementary Fig. 3**). Thus, our results add AMPAR surface diffusion to the variety of pre- and post-synaptic mechanisms that tune short-term synaptic depression of transmission between layers 4 and 2/3 in primary cortices, where it may contribute to frequency-dependent adaptation during prolonged sensory stimulation.

Many reports have demonstrated that AMPAR lateral diffusion can be tuned by neuronal activity ^4,7,13,21,22,26–28,30^. Our study now provides direct evidence that decrease in AMPAR mobility upon LTP induction regulates use-dependent synaptic transmission. Specifically, we used induction protocols to elicit LTP as a physiological means to modify AMPAR mobility and then measured its impact on synaptic transmission during high-frequency pre-synaptic firing. Utilizing post-synaptic photoactivatable CaMKII and either imaging receptor dynamics at synapses (**Fig. 3H**), or measuring synaptic responses to train stimuli (**Fig. 3I, K**), we demonstrate that purely post-synaptic CaMKII activity is sufficient to modify receptor diffusion-trapping and tune HF-STP. These findings are consistent with earlier observations that the changes in HF-STP after LTP induction are dependent on the post-synaptic activity of CaMKII ^46,52^. The transient increased HF-STP depression during eLTP could act as a negative feedback mechanism to prevent synapse over potentiation during consecutive trains of stimulation. Since the effects of the artificial NA cross-link we used in **Fig. 1 and 2** largely resemble those obtained at synapses potentiated by CaMKII activation (**Fig. 3E-K**), this promotes the NA cross-link approach as a tool with prospects to uncover implications of AMPAR diffusion-trapping mechanisms and short-term synaptic plasticity in brain function both *in vitro* and *in vivo*.

Finally, we also show that the effect of NA-mediated cross-link of AMPARs specifically mimicked enhancement of short-term synaptic depression that occurred transiently after LTP induction via electrical stimulation of L4 – L2/3 PN synapses in primary somatosensory cortex (**Fig. 3A-D**). This is consistent with studies showing a transient depressing HF-STP during early phases of LTP ^46^. Interestingly, the transient nature of LTP-induced HF-STP depression is in line with existing models that describe the trafficking of AMPARs during LTP, where diffusion-trapping of AMPARs play a vital role in the initial stages of LTP. However, as LTP progresses, the supply of extrasynaptic AMPARs is facilitated through the exocytosis of newly synthesized or recycled receptors which are highly mobile ^26,30^ and re-equilibrate the respective pools of mobile and immobile AMPARs in potentiated synapses. More generally, molecules that regulate AMPAR mobility could impact HF-STP. For example, experimentally manipulating the integrity of the extracellular matrix enhances AMPAR surface diffusion and short-term synaptic depression ^5^.

One of the effects concomitant with AMPAR immobilization-induced depression of post-synaptic responses during bursts of pre-synaptic firing (**Fig. 4A-D**) was a lower frequency of spontaneous network activity (**Fig. 4E-G**). The observed network effect associated with reduced summation of synaptic responses at high-frequencies is consistent with a putative role of synaptic depression in gain control - to dampen the strength of synapses during ongoing, repetitive pre-synaptic activity. The extent of gain control observed could also have been enhanced by the selective enhancement of short term-depression only at excitatory inputs, effectively acting then to reduce excitatory/inhibitory balance and dampen neuronal network activity as firing frequency increases ^53^. Another effect on spontaneous neuronal network observed after AMPAR immobilization-induced depression of burst responses was a lower coherence of neuronal firing, as measured by reduction in the pairwise correlation of neuronal firing across the network of neurons in hippocampal slice cultures (**Fig. 4H-I**). This effect is in-keeping with modelling studies that indicate enhanced propagation of synchronous neuronal firing in networks where synaptic connections are endowed with the capacity to undergo short-term facilitation ^54^. Since AMPAR immobilization was also associated with fewer neurons being recruited during spontaneous network activity, this opens the possibility that AMPAR immobilization-induced changes in HF-STP could lead to deterioration of engram-to-engram cell connections and compromise reactivation of engram cells in some contexts relating to memory retrieval ^55^.

Overall, our study reveals that AMPAR surface mobility is an intrinsic post-synaptic factor controlling HF-STP, in complement with glutamate release probability and AMPAR desensitization kinetics. In addition, both experimentally and physiologically instigated changes to the mobility of AMPARs controls the temporal integration of synaptic inputs. These data establish proof-of-concept for the role of AMPAR surface diffusion in regulating the activity of neural circuits. The recently developed *in vivo* tools used here to effectively mimic diffusional trapping of AMPARs by physiologically-relevant stimuli will be indispensable for evaluating AMPAR mobility and short-term dynamics of synaptic transmission on information processing in specific brain areas. The wealth of potential physiological regulators of AMPAR mobility, including LTP (this study) and LTD induction, hormones, signaling pathways or extracellular matrix elements ^15^ provides a previously uncovered diversity of post-synaptic mechanisms to control synaptic integration and network activity.

## Acknowledgments

We thank E. Verdier, N. Retailleau, N. Chevrier, and R. Sterling for cell culture and molecular biology support; H. El Oussini-Ben Chaabane, T.A. Vernoy, G. Dabee, P. Costet, C. Martin for support with animal husbandry; F. Lanore for support with GCaMP analysis; D. Rusakov, T. Jensen and S. Rama for support with iGluSnFR experiments. We thank the cell biology, PIV-EOPS and Plateforme In Vivo Experimental (PIV-EXPE) facilities of the IINS; the biochemistry and biophysics platform and animal genotyping facility of Neurocentre Magendie (INSERM); and the AAV production platform of the IMN. The microscopy was done at the Bordeaux Imaging Center, a service unit of CNRS-INSERM and Bordeaux University, a member of the national infrastructure France BioImaging.

## Authors contributions

Experimental conception and design: AN, AMG, DC. Manuscript preparation: AN, AMG, ACP, DC. Electrophysiology: AN, AMG, ACP, DBN. LLSM, 2-photon imaging & analysis: AMG, SM, MD. Molecular biology and biochemistry: SD, CB. In situ labeling: AN. dSTORM: HZ. uPAINT: HZ, AN. Animal surgeries: AN, AMG. Conception and production of mSA-ALFA: MS, CL.

## Data and materials availability

All data are available in the manuscript or the supplementary materials.

## Methods

### Plasmids

The BirA^ER^ coding sequence was a gift from A. Ting (*61*). The BirA^ER^-eGFP and BirA^ER^-Cre constructs were prepared as described in (*31*). For AP-SEP-GluA2 LY-BirA^ER^: mutagenesis on the GluA2 sequence was performed to encode the L483Y amino substitution. A fragment of the mutagenized GluA2 sequence between the BstEII/BspEI restriction sites was then subcloned into the AP-SEP-GluA2 construct (prepared as described in (*30*)) on a pBI-Tet on vector backbone (Clontech; #6152-1). For AP-SEP-GluA2 ET/YR-BirA^ER^: the sequence of the mutated GluA2 E713T Y768T was a gift from A. Plested (*41*). The sequence was subcloned to the AP-SEP-GluA2 construct on a pBI-Tet on vector backbone using XhoI/BspEI restriction sites. For BirA^ER^-AP-tdTomato-GluA1 (AP-tdTom-GluA1): the ORF of BirA^ER^ and AP-SEP-GluA1 were cloned into the lentiviral vector Syn-DsRed-Syn-eGFP (*62*) containing dual promoters by replacing respectively the DsRed and eGFP cDNA. Then the AgeI/NheI SEP cassette on GluA1 was replaced by tdTomato sequence. For AP-mRFP-GluA1: the SEP sequence in AP-SEP-GluA1 was replaced by the mRFP sequence using AgeI/NheI restriction sites on a pRK5 backbone. For the FLEx-mScarlet-P2A-paCaMKII: the mScarlet*-*P2A-paCaMKII sequence, which was a gift from H. Murakoshi (*52*), was amplified by PCR and inserted into the Cre-dependent AAV hSyn FLEx vector using BamHI/KpnI restriction sites.pTet-on (transactivator) was purchased from Clontech (Clontech; #P3070-5). Plasmids were prepared using the ZymoPURE Plasmid MaxiPrep Kit (Zymo Research, #ZD4203). All constructs were verified by restriction enzyme digestion and Sanger DNA sequencing. The amino acid position of mutations in GluA2 are defined with respect to amino acid numbering in the mature protein of the reference sequence (without the signal peptide of 21 amino acids).

AAV serotype 1 and 9 preps were produced by the viral core facilities of the Bordeaux Neurocampus IMN, Charité Universitätsmedizin Berlin, or ordered from Addgene. pAAV_synP.DIO.EGFP. WPRE.hGH (FLEx eGFP) was a gift from I. Wickersham (Addgene plasmid no. 100043; Addgene viral prep no. 100043-AAV9; https://www.addgene.org/100043; RRID:Addgene_100043). pAAV_hSyn.DIO.mCherry (FLEx mCherry) was a gift from B. Roth (Addgene plasmid #50459; http://n2t.net/addgene:50459; RRID: Addgene_50459). pAAV_hSynap.FLEX.SF.iGluSnFR. A184V (FLEx iGluSnFR) was a gift from L. Looger (*39*) (Addgene plasmid no. 106181; RRID: Addgene_106181). pGP-AAV_syn.FLEX.jGCaMP8m.WPRE (FLEx jGCaMP8m) was a gift from GENIE Project (Addgene plasmid #162378; http://n2t.net/addgene:162378; RRID: Addgene_162378) (*63*). pENN.AAV_hSyn.Cre.WPRE.hGH was a gift from James M. Wilson (Addgene plasmid #105553 ; http://n2t.net/addgene:105553 ; RRID:Addgene_105553). Viral titers were between 4.3 × 10^12^ and 8.3 × 10^13^ genome-containing particles (GCP)/ml.

### Animals

The AP-GluA2 knock in (KI) mouse line was generated as described in Getz et al. 2022 (*31*). AP-GluA2 KI mice used for slice culture and acute slice experiments were bred on a C57Bl6/J background. AP-GluA2 KI animals were obtained from heterozygote or homozygote mating and identified by genotyping. Genotyping was performed by polymerase chain reaction (PCR) assay on tail biopsies by the genotyping facility of Neurocentre Magendie (Bordeaux Neurocampus). Sprague-Dawley rats used for preparation of primary hippocampal cultures and slice cultures, were bred in the Plateforme In Vivo (PIV) facility of the Interdisciplinary Institute for Neuroscience (IINS). Animals were housed in 12/12 light/dark with *ad libitum* feeding. Every effort was made to minimize the number of animals used and their suffering. The experimental designs and all procedures were in accordance with the European guide for the care and use of laboratory animals and the animal care guidelines issued by the animal experimental committee of Bordeaux Universities (CE50; animal facilities authorizations A33063941 and A3306940; ethical project authorizations 20778-2019021913051936, 21848-2019083112401921, 38390-2022082913218868).

### Heterologous cell culture

HEK293T cells (ECACC, #12022001) were cultured at 37°C under 5% CO_2_ in DMEM (Gibco, #11995-065) supplemented with 10% FBS (fetal bovine serum), 1% L-glutamine (Thermo Fisher, #5030081) and 1% penicillin/streptomycin (Thermo Fisher, #15140-122).

### Primary rat hippocampal dissociated neurons

Primary dissociated hippocampal neuron cultures were prepared from pregnant rat females bred in the PIV facility of the IINS. Animals were handled and euthanized according to European ethical rules and protocols approved by the local ethics committee office 50. Dissociated hippocampal neurons from embryonic day 18 (E18) Sprague-Dawley rat embryos of either sex were prepared as described in (*64*). Briefly, dissociated neurons were plated at a density of 250,000 cells per 60 mm dish on 0.1 mg.mL-1 PLL pre-coated 1.5 h, 18 mm coverslips (Marienfeld Superior, #0117580). Neuronal cultures were maintained in Neurobasal™ Plus Medium (Thermo Fisher Scientific) supplemented with 0.5 mM GlutaMAX (Thermo Fisher Scientific; #35050061) and 1X B-27™ Plus Supplement (Gibco; #A3582801). 2 μM Cytosine β-D-arabinofuranoside (Sigma Aldrich, #C1768) was added after 72 h. At DIV7, cells were transfected with the respective cDNAs using a calcium phosphate procedure described in (*65*). For the super resolution NA cross-link experiments cells were transfected with AP-mRFP1-GluA1, AP-SEP-GluA2-BirA^ER^ and the Tet-on transactivatior and cultured in the presence of 10 μM biotin (Sigma-Aldrich; #B4501). 100 nM doxycycline (Sigma-Aldrich; #D1822) was added to the culture medium for 24h prior to the experiment. Astrocyte feeder layers were prepared from rat embryos, plated between 20,000 to 40,000 cells per 60 mm dish and cultured in Minimum Essential Medium (Thermo Fisher Scientific) containing 4.5 g.L^-1^ glucose, 2 mM GlutaMAX and 10% heat-inactivated horse serum for 14 days.

### Organotypic hippocampal slice preparation, transfection and transduction

Organotypic hippocampal slices were prepared and cultured according to the Muller interface method (*66*). At postnatal day 5 – 8 animals of either sex were anesthetized on ice and decapitated. Hippocampi were dissected from AP-GluA2 KI mice in ice-cold dissection medium composed of (in mM): 10 D-glucose, 4 KCl, 26 NaHCO_3_, 233.17 sucrose, 5 MgCl_2_, 1 CaCl_2_, 0.1% phenol red equilibrated with carbogen (95% O_2_, 5% CO_2_). Hippocampi from Sprague-Dawley rats were dissected in a sucrose-modified Gey’s balanced salt solution (in mM): 175 sucrose, 50 NaCl, 2.5 KCl, 0.85 Na_2_HPO_4_, 0.66 KH_2_PO_4_, 2.7 NaHCO_3_, 0.28 MgSO_4_, 2 MgCl_2_, 0.5 CaCl_2_, 25 glucose and 10 g/ml phenol red (330 mOsm/L, pH 7.3). Transverse 300 μm slices were cut with a tissue chopper (McIlwain), and transferred into ice-cold culture medium composed of: MEM powder (Sigma-Aldrich, #M4642), 20% inactivated horse serum (Gibco, #26050088), 1 mM L-glutamine (Thermo Fisher, #25030081), 1 mM CaCl_2_, 2 mM MgSO_4_, 1 mg/L insulin (Sigma, #I0516), 0.25 mM sodium L-ascorbate, 30 mM HEPES, 13 mM D-glucose, 5.2 mM NaHCO_3_ (pH 7.3, 320 mOsm/L). For rat OHS the culture medium consisted of: 50% Basal Medium Eagles (BME; Gibco; #21010046), 25% Hank’s balanced salt solution (HBSS; Gibco, #14025092), 25% inactivated horse serum (Gibco, #26050088), 1 mM L-Glutamine (Thermo Fisher, #25030081), and 6.5 g/L D-glucose (320 mOsm/L). Slices were then plated on ø 4 mm membrane segments (Millipore, #FHLC01300) placed on culture inserts (Millipore, #PICM0RG50) in 6-well plates containing 1 ml slice culture medium per well. Slices were maintained at 35 °C with 5% CO_2_ and the culture medium was replaced every 2 - 3 days.

CA1 PNs in mouse and rat organotypic hippocampal slices were transfected with single-cell electroporation (SCE) (*67*). Neurons in mouse OHS were transfected with BirA^ER^-eGFP. To overexpress AMPAR desensitization mutants in rat OHS, neurons were co-transfected with AP-SEP-GluA2 ET/YR-BirA^ER^ or AP-SEP-GluA2 LY-BirA^ER^, AP-tdTomato-GluA1 and the pTet-on transactivator. Briefly, on DIV3 slices were individually transferred to the chamber of an upright microscope (Nikon, Eclipse FN1). Before starting SCE, the microscope chamber was washed with 70% ethanol and allowed to dry. Slices were immersed in warm Tyrode’s solution composed of (in mM): 10 D-glucose, 10 HEPES, 120 NaCl, 3.5 KCl, 2 CaCl_2_, 2 MgCl_2_, 2 NaHCO_3_, 1 sodium pyruvate (Thermo Fisher, #11360070); (pH 7.3, 300 mOsm/L). Patch pipettes (approximately 4-6 MΩ) pulled from glass capillaries (Science Products GmbH, #GB150F-10P) were filled with potassium-based solution containing (in mM): 135 K-gluconate, 0.2 EGTA, 10 HEPES, 4 MgCl_2_, 4 Na_2_-ATP, 0.4 Na-GTP, 10 Na_2_-phosphocreatine, 3 L-ascorbic acid (pH 7.2, 290 mOsm/L) supplemented with plasmid DNA (10-13 ng/μl). After obtaining loose-patch seals, electroporation was achieved by delivering 4 x 25 ms pulses of 2.5 V at 1 Hz using a constant voltage stimulus isolator (NPI ISO-STIM 01D, Multi-Channel Systems STG 4002, Voltcraft FPS-1132). Upon returning the slices to the incubator the culture medium was supplemented with 10 μM D-biotin (for NA cross-link or NA/mSA labelling experiments) until experiments were performed. 10-20 cells in the CA1 per slice were electroporated with a success rate of approximately 50%.

Slices were transduced with: BirA^ER^-Cre + FLEx eGFP, Cre + FLEx eGFP, BirA^ER^-Cre + FLEx mCherry, BirA^ER^-Cre + FLEx jGCaMP8m, BirA^ER^-Cre + FLEx iGluSnFR and BirA^ER^-Cre + FLEx paCaMKII AAV9 at DIV1 by microinjection of the virus (es) (three to ten pulses, 30 ms, 69 kPa) into the CA1 pyramidal cell layer with glass microelectrodes (∼1 to 2 MΩ; Science Products, #GB150F-10P) (*68*). FLEx eGFP, FLEx mCherry, FLEx jGCaMP8m, FLEx iGluSnFR and FLEx paCaMKII viruses were used at a dilution of 1:10 – 1:20, BirA^ER^-Cre at 1:5000 to 1:20,000, Cre at 1:1000 in 1 x PBS. During the procedure, samples were maintained in medium on culture inserts and visualized under a stereomicroscope (Nikon SMZ 745T, Lumenera Infinity1). The microelectrode was positioned with a micromanipulator (Scientifica PatchStar), and viruses were injected using a Picospritzer (Parker Picospritzer III). Upon returning the slices to the incubator the culture medium was supplemented with 10 μM D-biotin (for NA cross-link or NA/mSA labelling experiments) until experiments were performed.

### Stereotaxic surgeries

Surgeries were performed on 5-6 week-old AP-GluA2 KI mice of either sex. The animals were fixed in a stereotaxic frame and kept under isoflurane anesthesia (4% induction, 1-2% surgery). Mice were injected subcutaneously with buprenorphine (0.1 mg/kg), 300 μl 0,9 % NaCl, and locally lidocaine (7 mg/kg). Thin glass pipettes pulled from Hirschmann® ringcaps® capillaries (Sigma, #Z611239), were filled with a mix of AAV9 BirA^ER^-Cre (6,7.10^13^ gcp/ml) diluted 1:5000 – 1:20000 in 0,9 % NaCl and FLEx eGFP (2,73.10^13^ gcp/ml) diluted 1:10 or BirA^ER^-Cre and FLEx iGluSnFR (2,1.10^13^ gcp/ml; 1:10). The viruses were injected into the CA1 area at coordinates in relation to Bregma (AP, -2 mm; ML, -1.6 mm; DV, -1.2 mm) or S1 (AP, -1.6 mm; ML, -3.3 mm; DV, -0.3 mm). For each injection site 250 nl of the virus mix was injected with a hand syringe. The pipette was then left in place for 5 min to allow diffusion of the viruses. Mice were left to recover on a heating pad for 2 – 3 h and then returned to home cages. The viruses were expressed for 2-3 weeks.

### Acute slice preparation

Acute slices from 8 – 10 week-old AP-GluA2 KI mice were prepared 3 weeks after the stereotaxic injections. Mice were anesthetized with a mix of Ketamine / Xylazine (130 mg and 13 mg/kg respectively) and cardiac-perfused with ice-cold sucrose-based solution composed of (in mM): 2 KCl, 26 NaHCO_3_, 1.15 NaH_2_PO_4_, 10 D-glucose, 220 sucrose, 0.2 CaCl_2_, 6 MgCl_2_ (320 mOsm/L). Brains were rapidly removed, dissected into 2 hemispheres, placed in the cutting chamber with ice-cold sucrose-based solution, and cut to 300 (or 275 μm for FRAP experiments) saggital (hippocampus) or coronal (cortex) slices with a Leica VT 1200S vibratome. Slices were transferred to the incubation chamber containing aSCF composed of (in mM): 126 NaCl, 2.5 KCl, 1.25 NaH_2_PO_4_, 25 NaHCO_3_, 12.1 D-glucose, 1.3 MgCl_2_, 2.5 CaCl_2_ (300 mOsm/L) placed in a water bath heated to 35°C. The water bath was then turned off and slices let to recover for 1h. All solutions were continuously equilibrated with carbogen (95% O_2_, 5% CO_2_).

### *In situ* labeling & wide-field microscopy

8 – 10 week-old AP-GluA2 KI mice expressing BirA^ER^-cre + FLEx eGFP in the CA1 or S1 were anesthetized with Ketamine / Xylazine (130 mg and 13 mg/kg respectively) and cardiac-perfused with PBS and 4% PFA. Brains were isolated and kept in 4% PFA o/n in 4°C. Brains were washed 2 x with PBS and left in 30% sucrose / PBS solution o/n in 4°C. Brains were washed in PBS and cut to 70 μm slices with a Leica VT 1200S vibratome. Slices were blocked with 2% Biotin-free BSA / PBS for 2h in RT, and incubated with 100 nM NeutrAvidin Protein DyLight 550 (Thermo Fischer, #84606) in 2% BSA biotin-free / PBS for 2h in RT. Slices were washed 2 x 10 min with 2% BSA biotin-free / PBS and 1 x 10 min with PBS. Next, they were incubated 10 min with 50 mM NH_4_Cl, washed 3 x with PBS and mounted on microscope slides with Fluoromount-G with DAPI (Southern Biotech; #C0822-P392). Samples were imaged with a DM5000 microscope (Leica) under HC PL Fluotar 10× numerical aperture 0.3 objective (Leica). Fluorescence excitation was obtained with a light-emitting diode SOLA light (Lumencor), and emission was captured by an ORCA-Flash4.0 V2 camera (Hamamatsu) controlled by MetaMorph software (Molecular Devices). Mosaics were done with a motorized XY stage (Märzhäuser).

### Super resolution microscopy

For dSTORM imaging, primary rat hippocampal neuronal cultures were treated with 50 nM NeutrAvidin (Thermo Fisher, #31000) in Tyrode’s solution composed of (in mM): 15 D-Glucose, 100 NaCl, 5 KCl, 2 MgCl2, 2 CaCl2, 10 HEPES (pH of 7.4, 247 mOsm/L) for 15 min to cross-link surface receptors. Then neurons were live labelled with monoclonal mouse anti-GluA2 antibody 15F1 (3,3µg/ml) (provided by E. Gouaux, Vollum Institute and Howard Hughes Medical Institute, Oregon Health and Science University, Portland, Oregon), at DIV16 for 7 minutes at 37° C. Cells were fixed for 10 min with 4% PFA-sucrose, washed 3 x with 1× PBS, incubated for 5 min with 50 mM NH_4_Cl, and washed 3 x with 1× PBS. Blocking was done for 30 min in 1× PBS with 2% BSA, and then cells were incubated for 30 min with goat anti-mouse IgG Alexa 647 (2µg/ml) (Thermo Fisher Scientific, #A21235) in 1× PBS with 2% BSA blocking solution. Cells were fixed with 2% PFA-sucrose, washed 3 x with 1× PBS, incubated for 5 min with 50 mM NH_4_Cl, and then washed 3 x with 1× PBS.

dSTORM imaging was performed on a LEICA DMi8 mounted on an anti-vibrational table (TMC, USA), using a Leica HCX PL APO 160x 1.43 NA oil immersion TIRF objective and fibber-coupled laser launch (405 nm, 488 nm, 532 nm, 561 nm and 642 nm) (Roper Scientific, Evry, France). Fluorescent signal was collected with a sensitive EMCCD camera (Evolve, Photometrics, USA). The 18 mm coverslips containing neurons were mounted on an open Ludin chamber (Life Imaging Services, Switzerland) and 600 µL of imaging buffer was added (*19*). To reduce oxygen exchange during acquisition an 18 mm coverslip was placed on top of the chamber. Image acquisition was driven by Metamorph software (Molecular devices, USA) and one image stack contained 40,000 frames. The size of the region acquired was 512×512 pixels (one pixel = 100 nm). Keeping the 642-laser intensity constant, the power of the 405 nm laser was increased during acquisition to control the level of single molecules per frame. Multicolor fluorescent microspheres (TetraSpeck, Invitrogen) were used as fiducial markers to register long-term acquisitions and correct for lateral drifts. Intensity-based drift-corrected super-resolution images with a pixel size of 25 nm were reconstructed, using PALMTracer software (*69-71*) operating as a plugin of MetaMorph. To determine the number of AMPARs per spine and analyse nanodomain properties, localization of Alexa-647 signals was performed using PALMTracer. AMPAR nanodomain properties were extracted from super-resolution images corrected for lateral drift as described in previous studies (*13, 19*) .

For the u-PAINT experiments surface AMPARs were cross-linked as described above. Neurons on 18 mm coverslip were placed onto a Ludin chamber (Life Imaging Services, Switzerland). The cells were kept in Tyrode solution at 37 °C. Imaging was done using a Nikon Ti-Eclipse microscope with an APO 100 × 1.49 NA oil immersion TIRF objective. Laser diodes with wavelengths of 405 nm, 488 nm, 561 nm, and 642 nm were used (Roper Scientific in Evry, France). To modify the illumination angle, a TIRF device (Ilas, Roper Scientific) was positioned in the laser path. The fluorescence signal was detected using a sensitive EMCCD camera (Evolve, Roper Scientific). The acquisition of images and control of the microscope were managed through Metamorph software (Molecular Devices, USA). The microscope was caged and heated to maintain the biological sample at 37°C. Neurons expressing mRFP-AP-GluA1 and SEP-AP-GluA2 were identified and low concentration of ATTO647-coupled GFP nanobody to reveal SEP-AP-GluA2 was added to the Ludin chamber. This allowed for sparse and stochastic labelling of endogenous GluA2-containing AMPARs at the cell surface. The TIRF angle was adjusted in an oblique configuration to detect the ATTO647 signal at the cell surface and minimize background noise from freely moving ATTO647-coupled nanobodies. A low-power 642 nm laser was activated to avoid photo-toxicity while maintaining a pointing accuracy of approximately 50 nm. Three movies, each consisting of 2000 frames at a rate of 33 Hz, were recorded to observe the surface diffusion of AMPARs. Single molecule localization, tracking and Mean Square Displacement (MSD) of ATTO647 signals (uPAINT) were computed using PALMTracer software like in Nair et al 2013 (*19*). From the MSD, the diffusion coefficient (D) corresponding to the global diffusion of the trajectory was calculated by linear fit of the first four points of the MSD plots.

### Electrophysiology in heterologous cells

cDNAs for pRK5_GluA1 (705.9 ng), pRK5_CMV.AP.TEV.GluA2 IRES.HA.BirAER (1176.5 ng), and pmaxGFP (117.6 ng) were co-transfected into HEK293T cells (60,000-62,500 cells.cm^-2^ in 6-well plate) using jetPRIME® (Polyplus-transfection, #114-01). 10 μM biotin and 40 μM NBQX were added to the cells at the time of the transfection. Cells were mechanically resuspended 1 day after transfection and seeded on PLL-coated coverslips in fresh media supplemented with biotin and NBQX. 2 days after transfection, cells were incubated in HBBS or 100 nM NA DyLight 550 (ThermoFisher, #84606) in HBSS for 20 min at 37°C. Cells were transferred to the recording chamber, and isolated GFP-positive cells were selected. NA-treated cells were selected based on GFP and DyLight 550 signal. Outside-out patch-clamp recordings were performed at room temperature in HEPES-buffered solution containing (in mM): 138 NaCl, 2 KCl, 2 MgCl_2_, 2 CaCl_2_, 10 D-glucose, and 10 HEPES (pH 7.4, 317-320 mOsm/L). Patch pipettes (resistance 4–6 MΩ) were filled with an internal solution containing (in mM): 120 cesium methanesulfonate, 2 NaCl, 2 MgCl_2_, 10 EGTA, 100 HEPES, and 4 Na_2_ATP (pH 7.4, 312 mOsm/L). Glu (10 mM) was dissolved in HEPES-buffered solution and applied using a theta pipette driven by a piezoelectric controller (Burleigh, #PZ-150M). Membrane potential was held at -60 mV. Currents were collected using an EPC10 amplifier (HEKA), filtered online and digitized at 20 kHz. Electrophysiological recordings were analysed with IGOR Pro 6 (WaveMetrics). Current amplitudes were measured with built-in tools, and τ_des_ was measured with exponential fit using a least-squares algorithm.

### Mouse organotypic slice electrophysiology

Electrophysiology experiments were performed 7-12 days after electroporation or 9 – 16 days after transduction (DIV10 – 17). For NA cross-link experiments, on the day of the experiment BirA^ER^-eGFP transfected slices were individually transferred to a 12-well dish, washed with 1 mL pre-warmed 1% BSA biotin free (Roth, #0132.2) / Tyrode’s solution (in mM: 10 D-glucose, 10 HEPES, 120 NaCl, 3.5 KCl, 2 CaCl_2_, 2 MgCl_2_, 2 NaHCO_3_, 1 sodium pyruvate (pH 7.3, 300 mOsm/L) and incubated in 1% BSA biotin free / Tyrode’s solution for 5 min. The slice was transferred onto a piece of parafilm and covered with a 30 μl droplet of 100 nM NeutrAvidin Protein (Thermo Fisher, #31000) in 1% BSA biotin free / Tyrode’s or 30 μl of 1% BSA biotin free / Tyrode’s alone (control) and incubated for 20 min. All incubation steps took place in 35 °C with 5% CO_2_. Next the slice was washed 2 x with 1 ml of pre-warmed 1% BSA biotin free / Tyrode’s and 1 x with pre-warmed Tyrode’s solution. Prior to electrophysiology recordings the slices were cut between CA3 and CA1 to prevent epilepsy. Slices were transferred to the microscope chamber perfused with carbogen equilibrated (95% O_2_, 5% CO_2_) aCSF composed of (in mM): 126 NaCl, 2.5 KCl, 1.25 NaH_2_PO_4_, 25 NaHCO_3_, 12.1 D-glucose, 1.3 or 4 MgCl_2_, 2.5 or 4 CaCl_2_, 0.002 gabazine (Tocris, #1262), 0.002 2-chloroadenosine (Tocris, #3136). High P_r_ conditions aCSF contained 1 mM MgCl_2_ and 4 mM CaCl_2_. Regular P_r_ conditions aCSF contained 1.3 mM MgCl_2_ and 2.5 mM CaCl_2_. The temperature of aCSF was kept at 30-34 °C by an in-line solution heater (WPI). A monopolar glass stimulation electrode, filled with aCSF and connected to a constant voltage stimulus isolator (IsoFlex, A.M.P.I.), was placed in the *stratum radiatum* of CA1. Electrode placement and patching was visually guided by observing the preparation with oblique illumination of infrared light through a 40 × water immersion objective (Nikon, NIR Apo 40× / 0.80W) and 1–2× optical zoom. Patch pipettes (4-6 MΩ) were pulled from borosilicate glass (Science Products GmbH, #GB150F-10P) and filled with a cesium-based internal solution composed of (in mM): 135 cesium methanesulfonate (Sigma-Aldrich, #C1426), 10 HEPES (Sigma-Aldrich, #H4034), 4 NaCl, 0.6 EGTA (Sigma-Aldrich, #3889), 0.1 CaCl_2_, 2 MgCl_2_, 2 Na_2_-ATP (Sigma-Aldrich, #A7699), 0.3 Na-GTP (Sigma-Aldrich, #51120), 3 QX314Cl (Tocris, #2313) and 0.15 spermine (Sima-Aldrich, #S4264) (pH 7.3, 290 mOsm/L). After obtaining a gigaseal, the cells were opened and maintained at -70 mV holding potential. Analogue voltage signals were filtered online and digitized at 10 kHz directly to the computer hard disk using an EPC-10 USB controlled by Patchmaster software (HEKA Electronik) from a computer workstation running Windows 10 (Microsoft). To record HF-STP, trains of 5 electrical stimulations (0.3 ms) at different frequencies (20 Hz, 10 Hz, 5 Hz, 2 Hz, 1 Hz) were delivered to the SC. The stimulation intensity was adjusted to obtain currents of 50 - 150 pA amplitudes. Each protocol consisted of 10 sweeps of trains delivered at 0.05 Hz. To calculate the EPSC_x_/EPSC_1_ ratio successful sweeps were averaged and the peak amplitude of consecutive EPSCs was measured using Clampfit 10 software (Molecular Devices). To quantify HF-STP profiles, the amplitudes of consecutive EPSCs in a train were normalized to amplitude of first EPSCs, summed and represented as cumulative normalized amplitude (CNA). The representative traces were scaled to the amplitude of EPSC_1_ in control condition.

For the AMPAR desensitization blocker experiments 50 μM D-APV (Tocris, #0106) and 200 μM CX546 (Tocris, #2980) were added to the aCSF. Paired recordings before and after the wash in of CX546 were performed. To measure mEPSCs, gabazine in the aCSF was replaced with 100 μM Picrotoxin (Tocris, #1128) and 0.5 μM Tetrodotoxin citrate (Tocris, #1069) was added to the aCSF. Recordings were performed at -70 mV holding potential. To measure the AMPAR rectification index, IV curves and AMPAR EPSC kinetics, ACSF was supplemented with 100 μM D-APV and 150 μM picrotoxin. To record IV curves 10 sweeps of single evoked AMPAR EPSC were recorded and averaged for each holding potential (-70, -60, -40, -20, 0, 20, 40 mV). A linear fit was performed to compute the theoretical EPSC amplitude at +40 mV. The rectification index was calculated as ratio between the theoretical and experimental value of AMPAR EPSC at +40 mV. For the recording of evoked NMDAR currents 150 μM Picrotoxin and 20 μM CNQX (Sigma-Aldrich, #C127) were added to the aCSF and cells were held at +40 mV.

For paCaMKII LTP experiments, OHS transduced with BirA^ER^-Cre + FLEx paCaMKII were cut between CA3 and CA1 and transferred to the microscope chamber perfused with equilibrated (95% O_2_, 5% CO_2_) aCSF composed of (in mM): 126 NaCl, 2.5 KCl, 1.25 NaH_2_PO_4_, 25 NaHCO_3_, 12.1 D-glucose, 1.3 MgCl_2_, 2.5 CaCl_2_, 0.002 2-chloroadenosine at 30 - 34°C. The internal solution was composed of (in mM): 135 cesium methanesulfonate, 10 HEPES, 4 NaCl, 0.6 EGTA, 0.1 CaCl_2_, 2 MgCl_2_, 2 Na_2_-ATP, 0.3 Na-GTP, 10 Na_2_-Phosphocreatine (Sigma-Aldrich, #P7936). mScarlet-paCaMKII positive neurons were patched. After reaching gigaseal configuration, cells were opened and held at -70 mV. A bipolar glass stimulation electrode, filled with aCSF, and connected to a constant current isolated stimulator (Digitimer Ltd.; #DS3), was placed in the *stratum radiatum* of CA1, and the amplitude of stimulation was adjusted to obtain currents of 50 – 150 pA. Electrode placement and patching was visually guided by observing the preparation with oblique illumination of infrared light through a 40× water immersion objective (Nikon, NIR Apo 40× /0.80W) and 1–2× optical zoom. The 3 min baseline of AMPAR EPSCs was elicited by stimulation at 0.033 Hz. During each sweep a test stimulation, and 2 seconds later, a HF-STP train of 5 stimulations at 20 Hz were delivered. Voltage signals were acquired and stored directly to the computer hard disk using an EPC-10 USB controlled by Patchmaster Next software (HEKA Electronik) from a computer workstation running Windows 10 (Microsoft). Photoactivation of mScarlet-paCaMKII consisted of 30 x 80 ms pulses at 1 Hz with 3 mW 460 nm, using a blue LED (*p*E-4000 CoolLED) triggered by Patchmaster. For sham control recordings, a minimal intensity of green light (550 nm) was triggered as above with the shutter closed. Cells were recorded for up to 30 min after photoactivation. To quantify early LTP/LTD profiles, the amplitudes of the first test stimulation 3 minutes before and after photoinduction were averaged and the outcome is presented as the ratio between the two. HF-STP profiles were measured as described above and the ratio of the average CNA 3 minutes before and after photoactivation is presented as a measure of the change in HF-STP after paCaMKII photoactivation.

### Rat organotypic slice electrophysiology

For experiments on GluA2 desensitization mutants, rat organotypic slices were electroporated with AP-tdTomato-GluA1, AP-SEP-GluA2-BirA^ER^ and rtTA transactivator on DIV7. On returning slices to the incubator, culture media was supplemented with 10 μg ml^−1^ gentamicin, 1 μg ml^−1^ doxycycline and 10 μM D-biotin until experiments were performed 2–3 weeks later. On the day of the experiment, organotypic slices were transferred to a storage chamber containing carbogen (95% O_2_ / 5% CO_2_)-bubbled aCSF (in mM): 125 NaCl, 25 NaHCO_3_, 1.25 NaH_2_PO_4_, 2.5 KCl, 2 CaCl_2_, 2 MgCl_2_, 10 D-glucose and 1 sodium pyruvate. A dialysis device (3,500 molecular weight cut-off, Slide-A-Lyzer MINI, Thermo Scientific) containing 0.25 ml of avidin Texas red conjugate was included in the storage chamber to quench free-biotin from the slices for at least 45 min. For the cross-link, slices were then individually transferred to a culture insert in a 35 mm culture dish containing 1 ml Tyrode’s solution with 100 nM NeutrAvidin and incubated at ambient temperature and atmospheric conditions for 45 min followed by washing in Tyrode’s solution before electrophysiology experiments. For the control, incubation was without NeutrAvidin (where the vehicle was Tyrode’s solution). Prior to any electrophysiological recordings of electrically evoked synaptic responses, CA3 was cut off to prevent seizures. For patch-clamp experiments, transfected slices were then transferred to the microscope chamber perfused with carbogen (95% O_2_ / 5% CO_2_)-bubbled recording aCSF maintained at approximately 30 - 34 °C by an in-line solution heater (WPI). The aCSF contained (in mM): 125 NaCl, NaHCO_3_, 2.5 KCl, 5 CaCl_2_, 1 MgSO_4_, 10 D-glucose, 1 sodium pyruvate, and 0.05 D-APV. For whole-cell patch clamp recordings, pipettes (∼5 MOhm) were filled with potassium-based intracellular solution containing (in mM): 135 K-methanesulfonate, 4 NaCl, 10 HEPES, 0.06 EGTA, 1 BAPTA, 0.01 CaCl_2_, 2 MgCl_2_, 2 Na_2_-ATP, 0.3 Na-GTP, 0.15 spermine, 0.05 Picrotoxin, (pH 7.3, 280 mOsm/L). Simultaneous field EPSP (fEPSP) recordings were made using patch pipettes filled with aCSF and positioned in *stratum radiatum* of CA1. For electrical field stimulation, a bipolar electrode fabricated from chlorided silver wires in a theta glass pipette with tip diameter (∼10 μm) filled with aCSF was positioned in *stratum radiatum* of CA1. All electrode placement and patching were visually guided by observing the preparation with oblique illumination of infrared light through a 40× water immersion objective (NIR Apo 40×/0.80W, Nikon) and 1–2x optical zoom. Stimulation (40 μs) intensity and polarity were adjusted using constant current stimulus isolator (IsoFlex, A.M.P.I.) until fEPSPs had an amplitude about ∼ 0.05 - 0.1 mV. The stimulus was delivered in a paired-pulse protocol with 9 sweeps with varying ISI (0.02, 0.06, 0.1, 0.18, 0.34, 0.66, 1.3, 2.58, and 5.14 s) and an intersweep interval of 20 s. Whole-cell recordings were made at holding potential of -80 mV without liquid junction potential correction or series resistance compensation. Analogue voltage signals were filtered online and digitized at 25 kHz directly to the computer hard disk either using an EPC-10 USB controlled by Patchmaster software (HEKA Electronik) from a computer workstation running Windows XP (Microsoft).

### Acute slice electrophysiology

Acute slices from AP-GluA2 KI mice were prepared 3 weeks after the stereotaxic injections as described above. Slices were recorded in a perfusion chamber continuously perfused with warm (30-34°C), carbogen equilibrated (95% O_2_ / 5% CO_2_) aCSF composed of (in mM): 126 NaCl, 2.5 KCl, 1.25 NaH_2_PO_4_, 25 NaHCO_3_, 12.1 D-glucose, 1.3 MgCl_2_, 2.5 CaCl_2_ (or 1 MgCl_2_ and 4 CaCl_2_ for high P_r_ conditions). A monopolar glass stimulation electrode, filled with aCSF, and connected to a constant voltage stimulus isolator (IsoFlex, A.M.P.I.), was placed in the *stratum radiatum* of CA1 (hippocampus) or L4 of S1 (cortex). Electrode placement and patching was visually guided by observing the preparation with oblique illumination of infrared light through a 40× water immersion objective (NIR Apo 40× /0.80W, Nikon) and 1 – 2× optical zoom. Patch pipettes (4-6 MOhm) were pulled from borosilicate glass (Science Products GmbH, #GB150F-10P) and filled with a cesium-based internal solution composed of (in mM): 135 cesium methanesulfonate, 10 HEPES, 4 NaCl, 0.6 EGTA, 0.1 CaCl_2_, 2 MgCl_2_, 2 Na_2_-ATP, 0.3 Na-GTP, 3 QX314Cl and 0.15 spermine (pH 7.3, 290 mOsm/L). After obtaining a gigaseal, the cells were opened and maintained at -70 mV holding potential. Analogue voltage signals were filtered online and digitized at 10 kHz directly to the computer hard disk using an EPC-10 USB controlled by Patchmaster software (HEKA Electronik) from a computer workstation running Windows 10 (Microsoft). To record HF-STP, trains of 5 electrical stimulations (0.3 ms) at different frequencies (20 Hz, 10 Hz, 5 Hz, 2 Hz, 1 Hz) were delivered. The stimulation intensity was adjusted to obtain currents of 50-150 pA amplitudes. Each protocol consisted of 10 sweeps of trains delivered at 0.05 Hz. To calculate the EPSC_x_/EPSC_1_ ratio, successful sweeps were averaged and the peak amplitude of consecutive EPSCs was measured using Clampfit 10 software (Molecular Devices). The amplitudes of consecutive EPSCs in a train were normalized to the amplitude of first EPSC, summed and represented as cumulative normalized amplitude. The representative traces were scaled to the amplitude of EPSC_1_ in control condition.

Somatic nucleated outside-out patches were pulled from CA1 and L2/3 pyramidal neurons in acute brain slices from 8 week-old AP-GluA2 KI mice. Slices were maintained in a perfusion chamber continuously perfused with warm (30-34°C), carbogen equilibrated (95% O_2_ / 5% CO_2_) aCSF composed of (in mM): 126 NaCl, 2.5 KCl, 1.25 NaH_2_PO_4_, 25 NaHCO_3_, 12.1 D-glucose, 1.3 MgCl_2_, 2.5 CaCl_2_. Patch pipettes (resistance 4–6 MΩ) were filled with an internal solution containing (in mM): 135 cesium methanesulfonate, 10 HEPES, 4 NaCl, 0.6 EGTA, 0.1 CaCl_2_, 2 MgCl_2_, 2 Na_2_-ATP, 0.3 Na-GTP, 3 QX314Cl and 0.15 spermine (pH 7.3, 290 mOsm/L). L-glutamate (1 mM) was dissolved in Ringer solution composed of (in mM): 135 NaCl, 5.4 KCl, 1.8 CaCl_2_, 1 MgCl_2_, 5 HEPES, 20 sucrose (pH 7.2, 300 mOsm/L) and applied using a theta pipette driven by a piezoelectric actuator and controller (PI; #P.245.50; #E-470.00). The other pipette barrel was filled with Ringer solution without sucrose. Membrane potential was held at -70 mV. Currents were collected using an EPC10 amplifier (HEKA), filtered online and digitized at 20 kHz. To measure the rate of desensitization a single 100 ms pulse of glutamate was applied. To measure recovery from desensitization, a single 100 ms pulse of glutamate was applied, followed by a second 100 ms pulse at increasing intervals (1 – 100 ms every 10 ms and 100 - 800 ms every 100 ms). τ_des_ was measured with a standard exponential fit. For τ_rec_, the amplitude of the current evoked by the second glutamate application was normalized to the amplitude of the first application current. τ_rec_ was calculated by one-phase association exponential fitting in Prism 8.2.1 (GraphPad).

For LTP experiments aCSF was supplemented with 150 μM picrotoxin and the internal solution was composed of (in mM): 135 cesium methanesulfonate, 10 HEPES, 4 NaCl, 0.6 EGTA, 0.1 CaCl_2_, 2 MgCl_2_, 2 Na_2_-ATP, 0.3 Na-GTP, 10 Na_2_-Phosphocreatine (Sigma-Aldrich, #P7936). To induce LTP in the L2/3 of the S1, the stimulation electrode was placed in the L4 and the amplitude of stimulation was adjusted to obtain currents of 50 – 150 pA. L2/3 PNs were patched and held at -70 mV. During the 5 min baseline a stimulation was delivered at 0.033 Hz. To induce LTP, the holding potential was switched to 0 mV and 50 stimulations at 0.167 Hz were delivered. In control conditions during the induction phase the holding potential was kept at -70 mV and the stimulation frequency at 0.033 Hz. Then, the holding potential was changed back to -70 mV and the AMPAR EPSC was recorded for another 30 min. After 5 min of baseline recording, directly after the end of LTP induction (or control) and every following 5 min, 4 sweeps of trains consisting of 5 pulses at 50 Hz were delivered (train 2 sec after the test stimulation) to measure HF-STP. To monitor the patch quality a test-pulse (5 mV, 10 ms) was delivered at the beginning of each sweep. Only recordings with R_s_<30 MΩ and where the R_s_ and R_m_ change throughout the recording was <25%, were analysed. To analyse HF-STP, each 4 sweeps with trains were averaged and analyzed as described above. For statistical analysis normalized AMPAR EPSC amplitude during last 10 min of recording was compared and analysed with an unpaired t-test.

### Photoactivation and FRAP with LLSM

The LLSM was built according to the technical information provided by the group of E. Betzig and used under license from Jamelia Research Campus, Howard Hughes Medical Institute (HHMI), USA (*72*). The LLSM was modified by addition of a photostimulation module (PSM) for one- or two-photon photomanipulation, as previously described in detail (*31*). The microscope was controlled by the Lattice Scope user interface written in LabVIEW (National Instruments). For the FRAP experiments following paCaMKII photoactivation, AP-GluA2 KI organotypic hippocampal slices were prepared and transduced with BirA^ER^-cre and FLEx-mScarlet-P2A-paCaMKII as described above. Slices were washed with 1 mL Tyrode’s solution (in mM: 10 D-glucose, 10 HEPES, 120 NaCl, 3.5 KCl, 2.5 CaCl_2_, 1.3 MgCl_2_, 2 NaHCO_3_, 1 sodium pyruvate, 1% BSA biotin free (pH 7.3, 300 mOsm/L)) for 5 min. Then slices were transferred onto parafilm and incubated for 20 min in 30 μl 1% BSA biotin free/Tyrode’s containing 400 nM monomeric StreptAvidin (mSA), which was engineered to contain 4 replicates of the ALFA tag on the N-terminus (*50, 73, 74*). Next, slices were washed twice with 1% BSA biotin free/ Tyrode’s and then incubated for 20 min with 400 nM ALFA nanobody coupled to Atto 643 (AD 643, Atto-Tec) (*50*). All incubation steps took place at 35 °C with 5% CO_2_. To confirm efficient NA penetration into OHS slices, slices were incubated with 100 nM NA conjugated to STAR 635P (Abberior, ST635P) for 20 min, with blocking and washing steps as above. Membranes were adhered onto 5 mm glass coverslips coated with poly-L-lysine and mounted onto the LLSM sample holder using Dow Corning high-vacuum silicone grease (Sigma-Aldrich, #Z273554). The imaging chamber was maintained at a temperature of 34°C and continuously perfused with aCSF (in mM: 12.1 D-glucose, 126 NaCl, 2.5 KCl, 1.3 MgCl_2_, 2.5 CaCl_2_, 25 NaHCO_3_, and 1.25 NaH_2_PO_4_ (300 mOsm/L), which was equilibrated with carbogen and perfused at a rate of 1.7 ml/min. Before starting the imaging experiments, slices were equilibrated for approximately 20-30 min and maintained in the imaging chamber for up to 2 hours. The power of illumination ranged from ∼20 to 200 µW spread over the entire width and thickness of the LLSM excitation plane, and was kept consistent across relevant experiments. Images were acquired at ∼10 to 30 µm beneath the surface of the slice. To identify ROIs and verify labelling specificity, 3D images (12.8 µm by 12.8 µm by 30 µm) of dendrites in CA1 *stratum radiatum* were acquired before FRAP imaging, with 559 nm excitation of mScarlet as a volume marker and BirA^ER^ or paCaMKII reporter, and 642 nm excitation of Atto 643 on the mSA-ALFA/ALFA-nb complex as the extracellular label for bAP-GluA2. 2-photon photoactivation of paCaMKII was 30 x 80 ms pulses at 1 Hz with 3 mW at 900 nm. Spine volume changes were followed with 3D acquisitions (25.6 µm by 25.6 µm by 40 µm) at 0.017 Hz during a 5 min baseline and for 3 min following photoactivation. FRAP illumination was 60 ms with 642 nm to a single focal point, the power measured at the back aperture of the detection objective was below 4 mW. Fluorescence recovery was followed by single-plane acquisitions (100 ms) in three steps at 10 Hz (15 s), 1 Hz (60 s), and 0.2 Hz (180 s). Calculation of the recovery curves and analysis of the recovery fraction were done as previously described (*31*). Quantification of bAP-GluA2 intensity over the surface of dendritic segments and spines and spine volumes were analysed on 3D images using the Filaments and Surfaces modules in Imaris 10.0.1 (Oxford Instruments), respectively. Intensity measurements were normalized to the mean intensity of a 3D background ROI of a standard size.

For acute slice imaging, slices from 8-10-week-old AP-GluA2 KI mice stereotaxically injected with BirA^ER^-cre + FLEx eGFP to the CA1 or S1, were transferred to a 24-well plate and incubated for 20 min in 0.5 ml aCSF with 400 nM mSA-ALFA, washed twice in aCSF, then incubated with 400 nM of the Atto 643 coupled ALFA nanobody. All incubation steps took place at room temperature with continuous carbogen equilibration (95% O_2_, 5% CO_2_). To confirm efficient NA penetration into *ex vivo* slices, slices were incubated with 100 nM NA conjugated to STAR 635P (Abberior, ST635P) for 20 min, with washing steps as above. We designed a modified LLSM sample holder with a harp and nylon grid to mount the slice on top of 5 mm poly-L-lysine coated glass coverslips. The modified sample holder and harp were designed with Autodesk Inventor Pro and manufactured of 316L stainless steel (Mecaprocess). 3D CAD files are available upon request. 3D images of dendrites in CA1 *stratum radiatum* or basal dendrites of L2/3 pyramidal neurons in S1 were acquired as above, with 488 nm excitation of eGFP as a volume marker and BirA^ER^ reporter, and 642 nm excitation of Atto 643 on the mSA-ALFA/ALFA-nb complex as the extracellular label for bAP-GluA2. Due to the drift observed in acute slice preparations, FRAP illumination was 30 ms to a single focal point and fluorescence recovery was followed using 3D acquisitions (12.8 µm by 12.8 µm by 10 µm, 30 ms) in three steps at 1 Hz (30 s), 0.5 Hz (100 s), and 0.2 Hz (150 s). Calculation of the recovery curves and analysis of the recovery fraction were done as above using maximum intensity projections that were corrected for drift using the BIG-EPFL StackReg plugin (Fiji, NIH). 3D quantification of bAP-GluA2 intensity was analyzed with Imaris, as above.

### 2-photon Biosensor Imaging

For iGluSnFR imaging experiments, organotypic hippocampal slices were prepared from AP-GluA2 KI mice and transduced with FLEx iGluSnFR and BirA^ER^-Cre as described above to verify biotinylation of AP-GluA2 in CA1 PNs (post-synaptic expression of the sensor). For jGCaMP8m imaging, slices were transduced with BirA^ER^-cre + FLEx jGCaMP8m as described above. SF-iGluSnFR-A184V and jGCaMP8m imaging was performed using a FEMTOSmart Femto2D dual scan-head 2-photon microscope (Femtonics) mounted on a BX61WI stand (Olympus) under a CFI Apo LWD 1.1 numerical aperture 25× 2.0-mm objective (Nikon). Chameleon Vision II or Ultra II lasers (Coherent) were used. For 2-photon excitation, laser power was modulated by 2 Pockel cells (Conoptics). Imaging was done through the galvo scanner or resonant scanner pathway. Fluorescence was collected on 2 upper and optionally on 2 lower NDD GaAsP photomultiplier tubes (PMTs) with green and red emission filters (Hamamatsu). The microscope was equipped with an electrophysiology system consisting of an Axon Instruments MultiClamp 700A amplifier and Digidata 1550A digitizer (Molecular Devices), a DS3 isolated current stimulator (Digitimer) and SM-5 micromanipulators (Luigs & Neumann). For 2-photon biosensor imaging, the microscope was controlled with MES (v 6.6.12344, Femtonics) software running in MATLAB (v 7.5.0.342, MathWorks) or MESc (v 4.0.2.11689, Femtonics) software and the electrophysiology system was controlled with the pCLAMP 10.7 software suite (Molecular Devices). The imaging chamber was maintained at 30-34°C with an in-line heater (TC-324C, Warner Instruments) and perfused with aCSF, which was equilibrated with carbogen and perfused at a rate of 1.7 ml/min. For high P_R_ iGluSnFR experiments aCSF contained (in mM): 12.1 D-glucose, 126 NaCl, 2.5 KCl, 1 MgCl_2_, 4 CaCl_2_, 25 NaHCO_3_, and 1.25 NaH_2_PO4 (300 mOsm/L). For normal P_R_ iGluSnFR and synaptic jGCaMP8m experiments aCSF contained instead (in mM): 1.3 MgCl_2_ and 2.5 CaCl_2_. For spontaneous jGCaMP8m experiments aCSF contained instead (in mM): 1.3 MgCl_2,_ 2.5 CaCl_2_ and 3.5 KCl. For NA cross-linking experiments, organotypic slices were incubated for 20 min in 1% BSA-Tyrode’s (as above) with or without 100 nM NA before being transferred into the imaging chamber, and acute slices were perfused in the imaging chamber with aCSF with or without 100 nM NA for 20 min. Slices were equilibrated in the chamber for approximately 20-30 min before imaging experiments and maintained in the imaging chamber for up to 2 hours. Illumination power ranged from ∼5 to10 mW. Line and area scans were done with the galvo scanner for 1.2 s at 500 Hz for iGluSnFR imaging or 4 s at 30 Hz for jGCaMP8m synaptic imaging, and HF-STP trains (5x or 10x at 20 Hz) were delivered using a bipolar glass stimulation electrode filled with aCSF and positioned ∼10-30 µm from the spine. Stimulation amplitude was typically 20 – 150 µA. 3-10 replicates per spine were acquired at a rate of ∼0.017 Hz and the spine ROI was manually repositioned between each acquisition to offset 3D drift. Peak analysis was done in Fiji (NIH) using custom macros. To quantify glutamate release profiles during HF-STP trains, the amplitudes of consecutive iGluSnFR peaks, normalized to the amplitude of the first peak, were summed and represented as cumulative normalized amplitude. To quantify spine calcium influx, the area under the curve and maximal response amplitudes were measured in spine ROIs. Response amplitudes were calculated as ΔF/F_0_ = ((F_peak_ − F_bkgd_) − (F_bsln_ − F_bkgd_)) / (F_bsln_ − F_bkgd_). For spontaneous jGCaMP8m imaging, 3x 60 s area scans per ROI were done with the resonant scanner at 100 Hz and the acquisitions were denoised by 3-fold averaging. ΔF/F computation, peak detection, frequency analysis and cross-correlation were done in MATLAB version 9.14.0 (R2023a Update 3; MathWorks Inc.) using the findpeaks and cross-correlation functions. Peak amplitudes were calculated as ΔF/F = ((F) − (F_bsln_)) / (F_bsln_)) where F_blsn_ was the 20^th^ percentile of cell ROI mean values.

### Statistical analysis

All reported results are derived from at least three independent experiments to ensure replicability. No statistical test was used to predetermine sample sizes. For imaging and electrophysiology data, all analysis was performed blinded to the experimental condition. Electrophysiology experiments involving NA cross-link were performed blind to the experimental condition. jGCaMP8m experiments were performed blind. Statistical analysis and data plotting were performed with Prism 8.2.1 or 9.4.0 (GraphPad). Datasets were analysed with the Shapiro-Wilk test for normality, and parametric (P > 0.05) or nonparametric (P < 0.05) statistical tests were performed as appropriate. F test or Bartlett’s test was used to assess equality of variance. Electrophysiological and super-resolution data was analysed with the robust regression and outlier removal (ROUT) method to identify outliers. Data is presented as Mean ± SEM unless otherwise stated. Test details and statistical outcomes for all experiments are reported in the relevant figure legends.

---

**Supplementary Figure 1.**
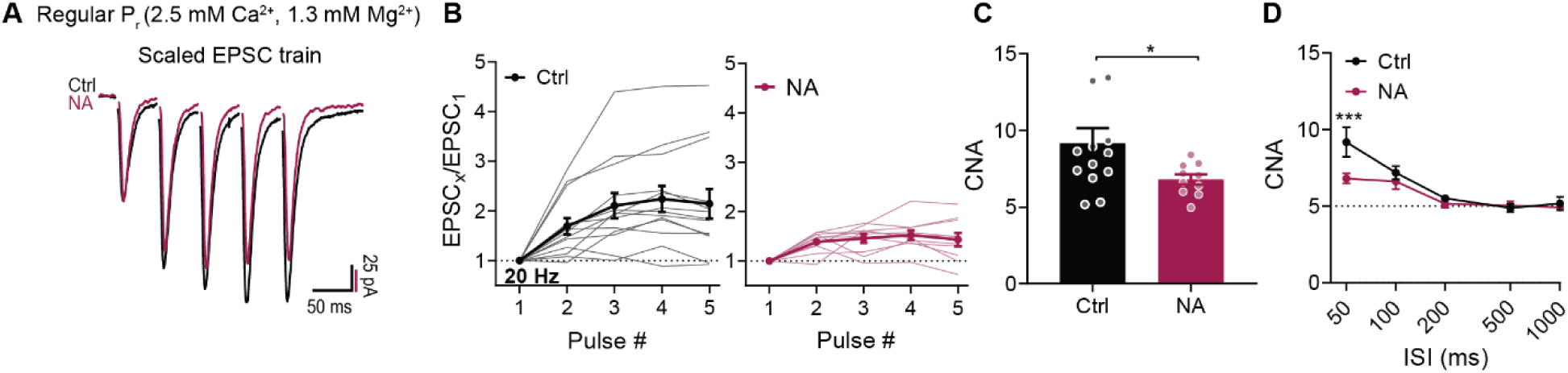
AMPAR surface diffusion regulates HF-STP in regular calcium conditions. (**A**) AMPAR EPSCs 20 Hz trains ± NA cross-link. (**B**) Whole-cell voltage clamp recordings of trains in SC – CA1 in OHS ± NA. Amplitude of AMPAR EPSC normalised to first response. Population average ± SEM in bold (n= 13, 10 cells). (**C**) Data from c represented as CNA (*P=0.035; Welch’s test). (**D**) CNA of trains ± NA with different ISI (n=11-13 and 10-12 cells; F=5.393, P=0.0221, ***P=0.0006; Two-way ANOVA, Fisher’s LSD post-hoc test).

**Supplementary Figure 2.**
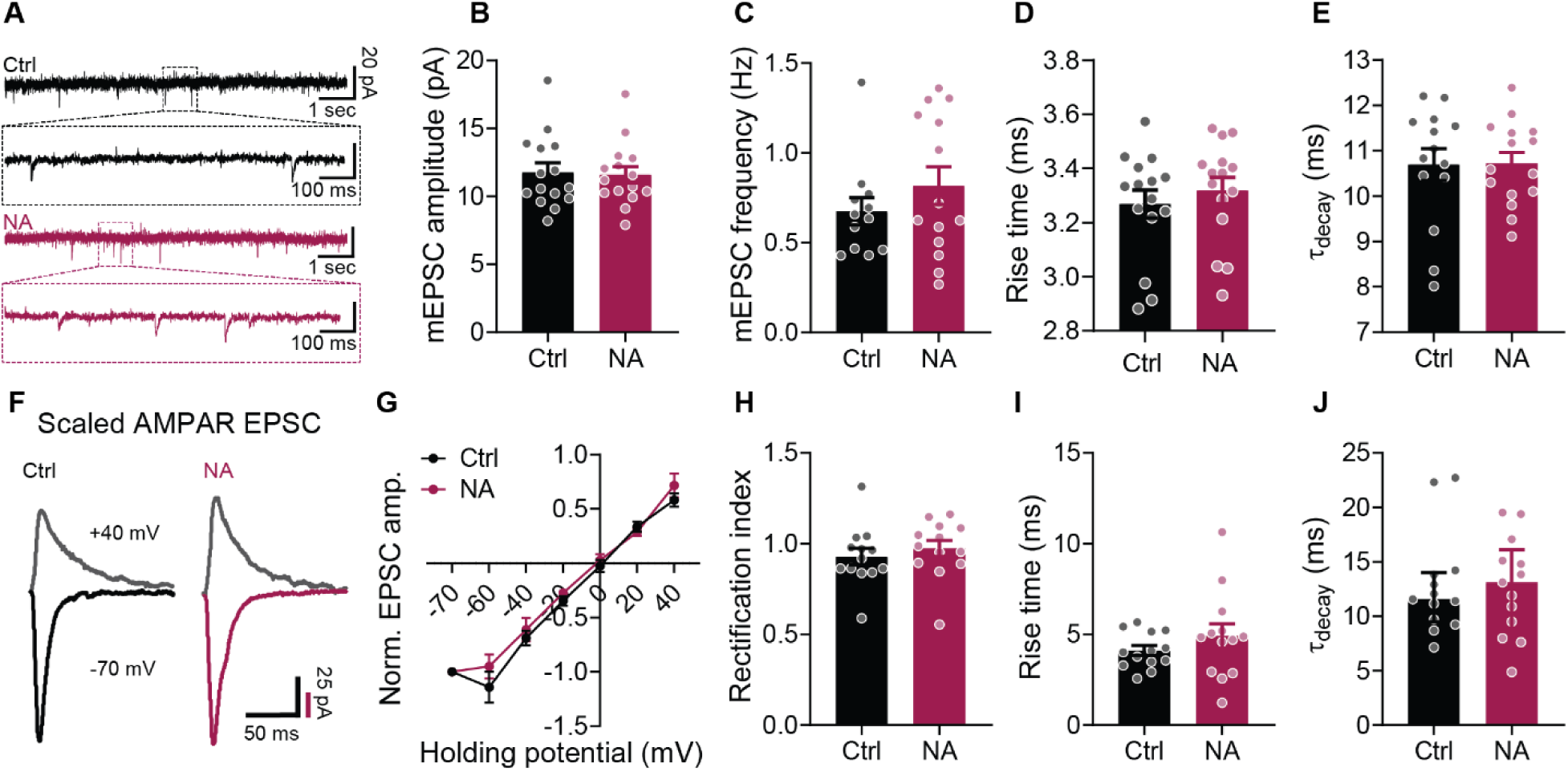
NA cross-link does not impact basal synaptic transmission. (**A**) mEPSCs recorded from CA1 PNs in OHS ± NA cross-link. (**B**) mEPSC amplitude ± NA ( n=15 cells; P=0.8409; Unpaired t-test), (**C**) frequency (n=12, 14 cells; P=0.6402; Mann-Whitney test), (**D**) rise time (10 – 90%); (n=15 cells; P=0.4901; Unpaired t-test), (**E**) τ of decay; (n=14, 15 cells; P=0.9473; Unpaired t-test). (**F**) Whole-cell voltage-clamp recordings of evoked AMPAR EPSCs in CA1 PNs in OHS ± NA cross-link: (**G**) I/V curves (normalised to amplitude at -70mV; n=13 cells), (**H**) rectification index (P=0.4960; Unpaired t-test), (**I**) rise time (10 – 90%); (P=0.2924; Welch’s test), (**J**) decay time (τ of decay); (Median ± quartiles; P=0.8010; Mann-Whitney test).

**Supplementary Figure 3.**
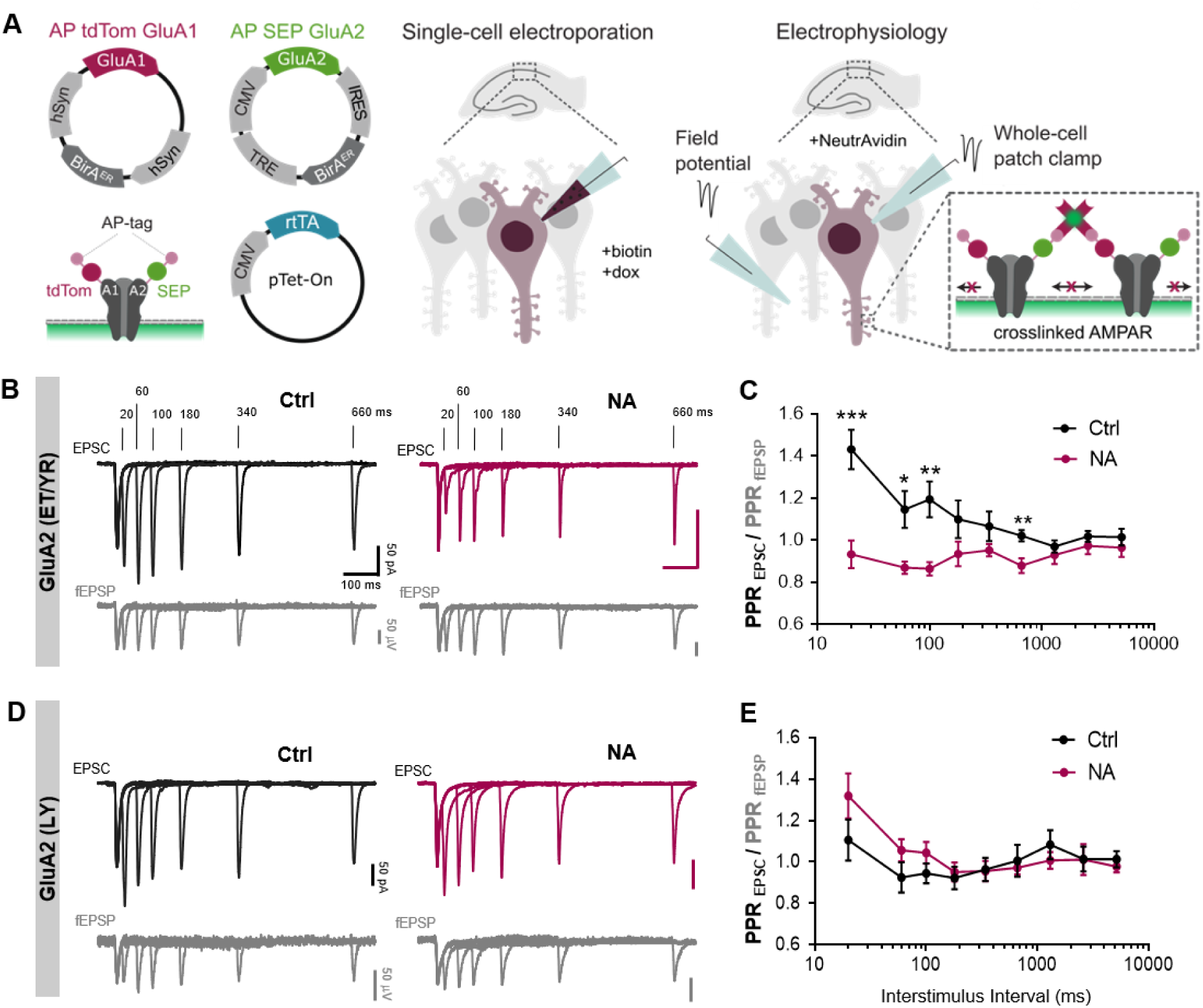
AMPAR desensitization mutants are more susceptible to immobilization-dependent HF-STP impairment. (**A**) Scheme of the experimental approach. Biotinylated tdTomato-GluA1 (AP-tdTom-GluA1), and Tet-on inducible biotinylated SEP-GluA2 (AP-SEP-GluA2) mutants to inhibit desensitization (L483Y, LY) and slow recovery from desensitization (E713T, and Y768R, ET/YR) were introduced to CA1 PNs by SCE. Whole-cell voltage clamp recordings of a paired pulse protocol with varying ISI were performed from CA1 PNs in OHS ±NA. To control for slice-to-slice variability in HF-STP, paired-pulse ratios of fEPSPs were simultaneously measured. aCSF was supplemented with 50 µM D-APV and with 1 and 5 mM of Mg^2+^ and Ca^2+^, respectively. (**B**) Exemplary recordings of EPSCs in neurons expressing the ET/YR mutant of biotinylated AP-SEP-GluA2. Below, matching exemplary control fEPSP. Traces shown are ensemble averages of multiple trials for each ISI. (**C**) Group data for the conditions shown in **B** (n= 8, 9 cells; F=13.79, P=0.0021, *-***P≤0.0160; Two-way ANOVA, Fisher’s LSD post hoc test). (**D**) Data representation as in **B** but for neurons expressing the LY mutant of biotinylated AP-SEP-GluA2. (**E**) Group data for the conditions shown in **D** (n= 12, 11 cells; F=0.5630, P=0.4614; Two-way ANOVA, Fisher’s LSD post hoc test).

**Supplementary Figure 4.**
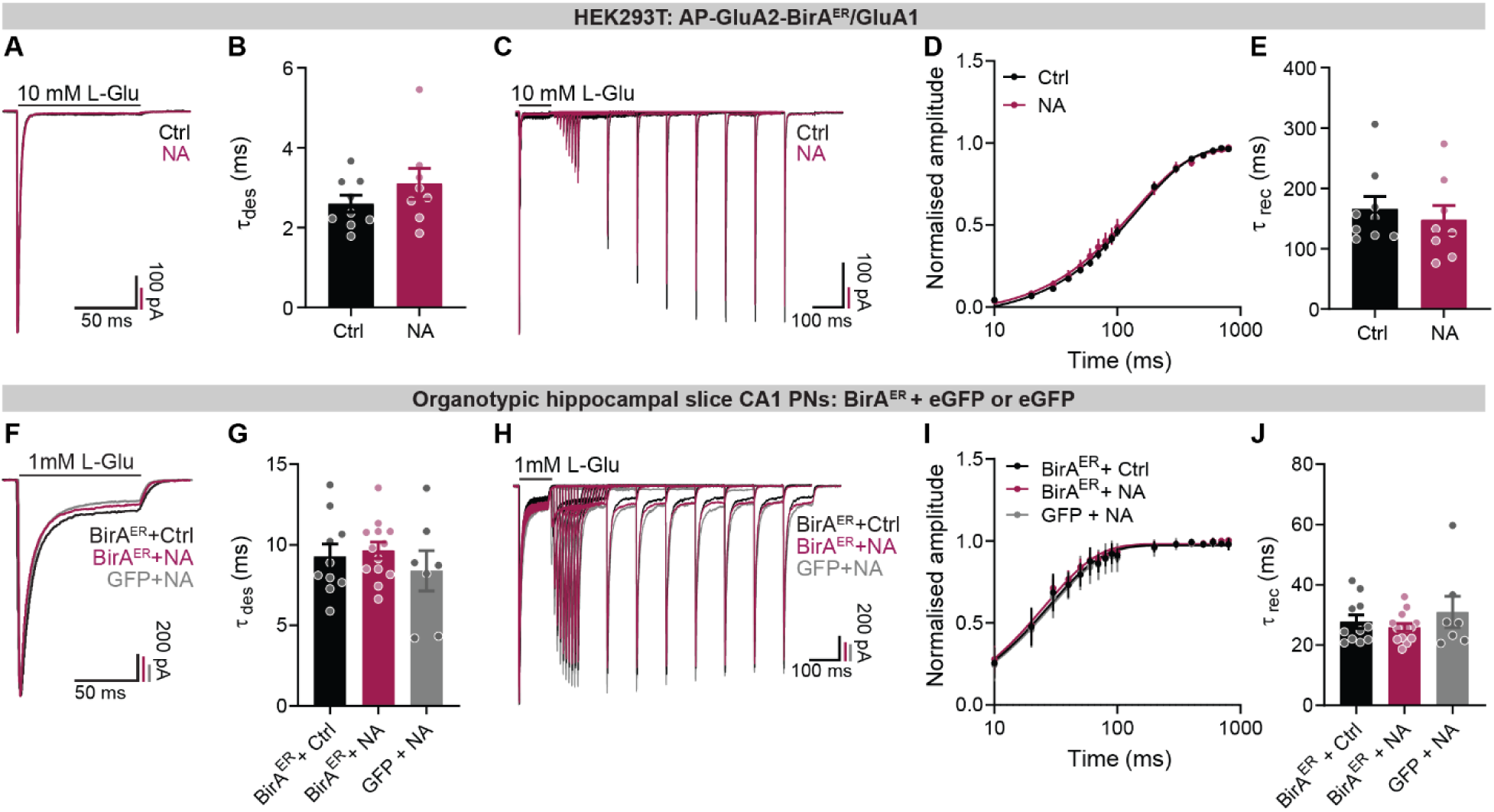
NA cross-link does not affect AMPAR kinetics. (**A – E**) Outside-out patch-clamp recordings with fast glutamate application from HEK293T cells expressing AP-GluA2/GluA1 and BirA^ER^ ± NA. (**A**) Exemplary traces of AMPAR current kinetics during 100 ms glutamate application. (**B**) τ of AMPAR desensitization (n=9, 8 cells; P=0.2585; Unpaired t-test). (**C**) AMPAR recovery from desensitization during 2-pulse glutamate application at increasing time intervals. (**D**) Averaged recovery from desensitization ± SEM and one-phase association exponential curve fitting (n=8, 8 cells). (**E**) τ of recovery from desensitization (P=0.9827; Unpaired t-test). Same experiment in OHS. Somatic nucleated outside-out patches excised from CA1 PNs expressing BirA^ER^ + eGFP or eGFP alone ± NA. (**F**) AMPAR current kinetics and (**G**) rate of desensitization (n =10, 13, 7 cells; F=0.6138, P=0.5487, P≥0.2768; One-way ANOVA, Fisher’s LSD post-hoc test). (**H - I**) Averaged recovery from desensitization ± SEM and one-phase association exponential curve fitting (n=11, 14, 7 cells). (**J**) τ of AMPAR recovery from desensitization (P=0.8510; Kruskal-Wallis test).

**Supplementary Figure 3.**
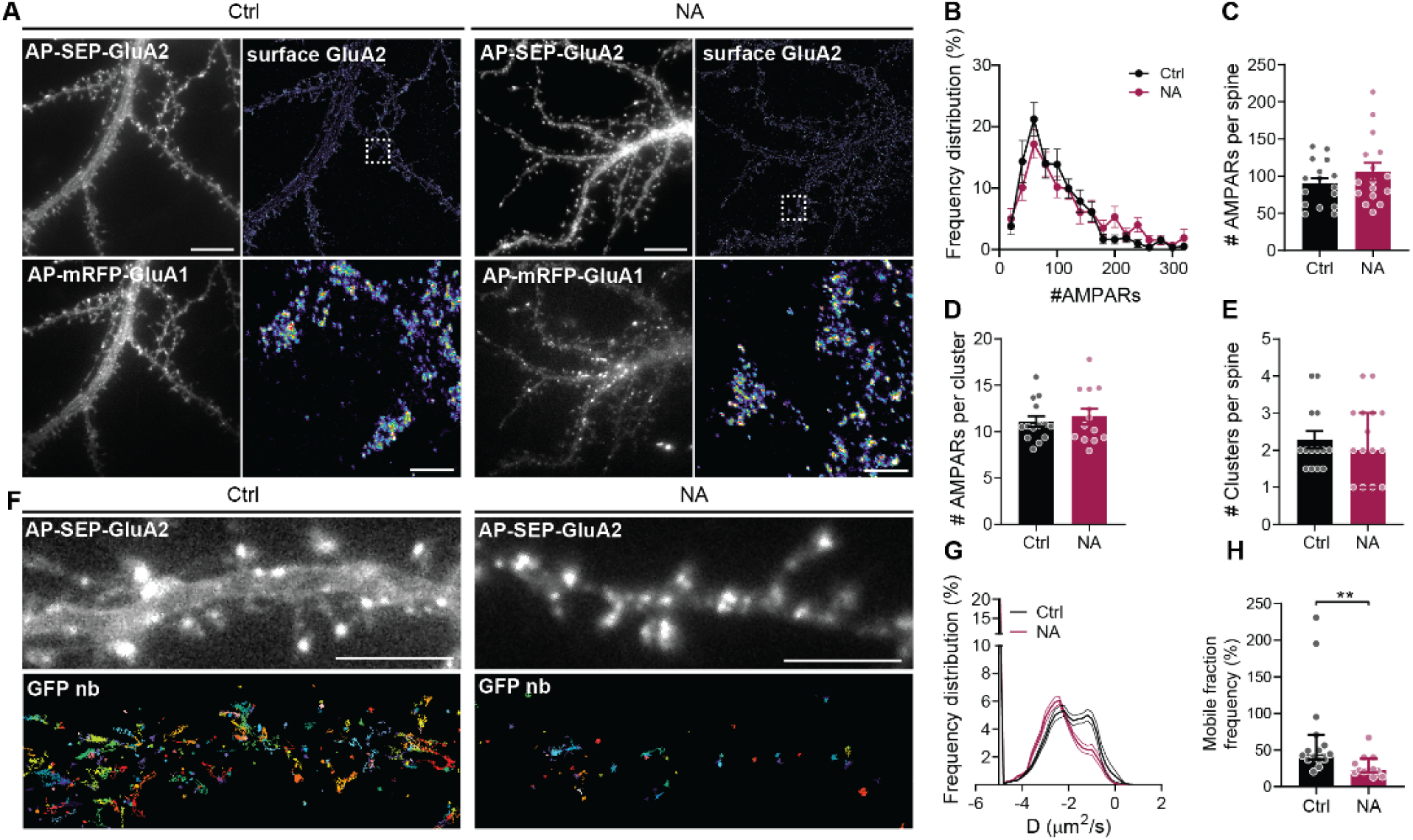
NA cross-link of AMPARs does not affect AMPAR nano-organization. (**A**) Wide-field (left) and super-resolution (right) images of neurons ± NA cross-link. Scale bar, 10 and 1 μm. (**B**) Frequency distribution of AMPAR number per spine (n=16 cells). (**C**) Number of GluA2-containing AMPARs per spine (n=16 cells; P=0.2429; Unpaired t-test), (**D**) GluA2-containing AMPARs per cluster (n=14, 13 cells; P=0.5843; Unpaired t-test), (**E**) clusters per spine (Median ± quartiles; n=14, 15 cells; P=0.9058; Mann-Whitney test). (**F**) uPAINT image acquisition. SEP-GluA2 signal (top) and GluA2-containing AMPAR tracks (bottom). Scale bar, 5 μm. (**G**) Frequency distribution of surface GluA2 diffusion rate (n=15, 12 cells). (**H**) Quantification of GluA2 mobile surface fraction ± NA (Median ± quartiles; n=15, 12 cells; **P=0.0017; Mann-Whitney test).

**Supplementary Figure 6.**
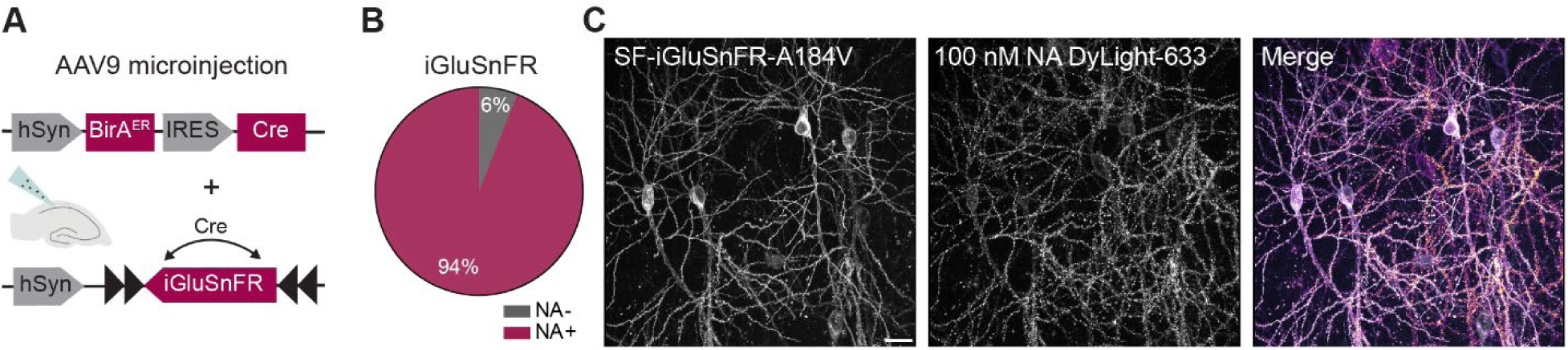
Verification of NA binding to bAP-GluA2. (**A**) Scheme of virus infection strategy to express SF-iGluSnFR-A184V and BirA^ER^ in CA1 PNs of OHS. (**B**) Percentage of iGluSnFR positive cells with detectable binding of NA to bAP-GluA2 (n=78/83 cells, 3 slices). (**C**) Confocal images of transduced CA1 PNs labeled with NA. Scale bar, 20 μm.

**Supplementary figure 7.**
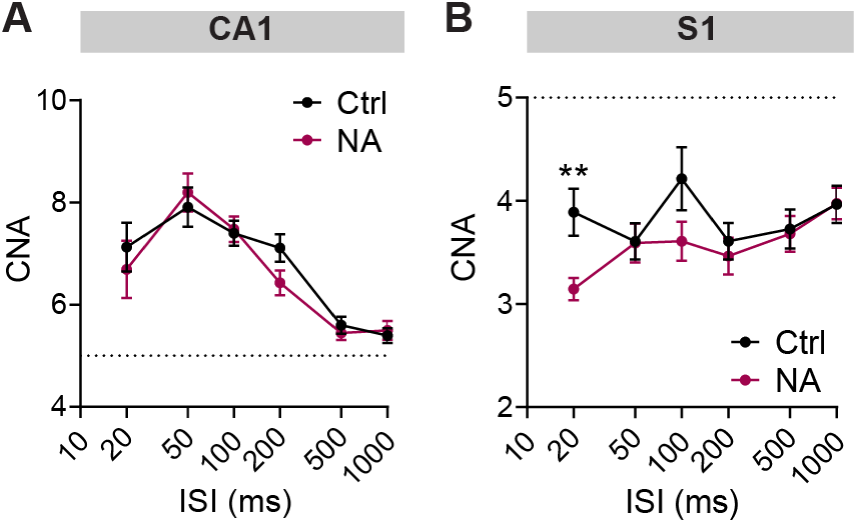
AMPAR cross-link enhances synaptic depression in S1, but not CA1, pyramidal neurons during high-frequency activity. (**A**) CNA of HF-STP trains induced with ISI of different duration in SC – CA1 (n=9-14 and 11 cells; F=0.4919, P=0.4845; Two-way ANOVA, Fisher’s LSD post-hoc test) and (**B**) L4-L2/3 of the S1 (n=16-19 and 17-20 cells; **P=0.0095; Mann-Whitney test).

**Supplementary Figure 8.**
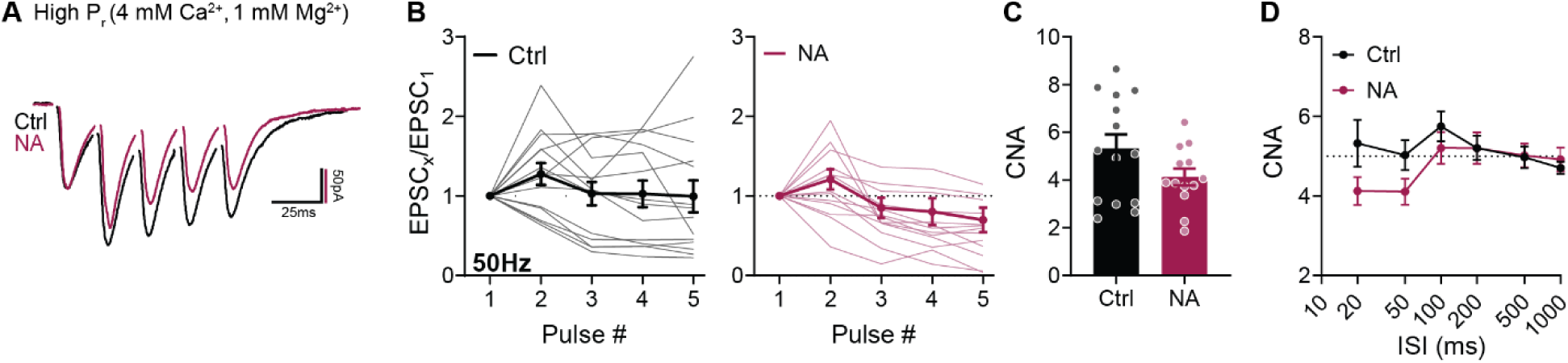
Effect of AMPAR mobility on HF-STP in the SC – CA1 synapse of *ex vivo* brain slices in high P_r_ conditions. (**A - D**) HF-STP 50 Hz trains recorded from CA1 PNs ± NA cross-link. (**B**) Population average ± SEM in bold (n=14, 13 cells). (**C**) Data from **B** represented as CNA of trains (P=0.0998; Unpaired t-test). (**D**) CNA of trains with different ISI (n=11-18 and 13-15 cells; P≥0.1227; Mann-Whitney test).

**Supplementary Figure 9.**
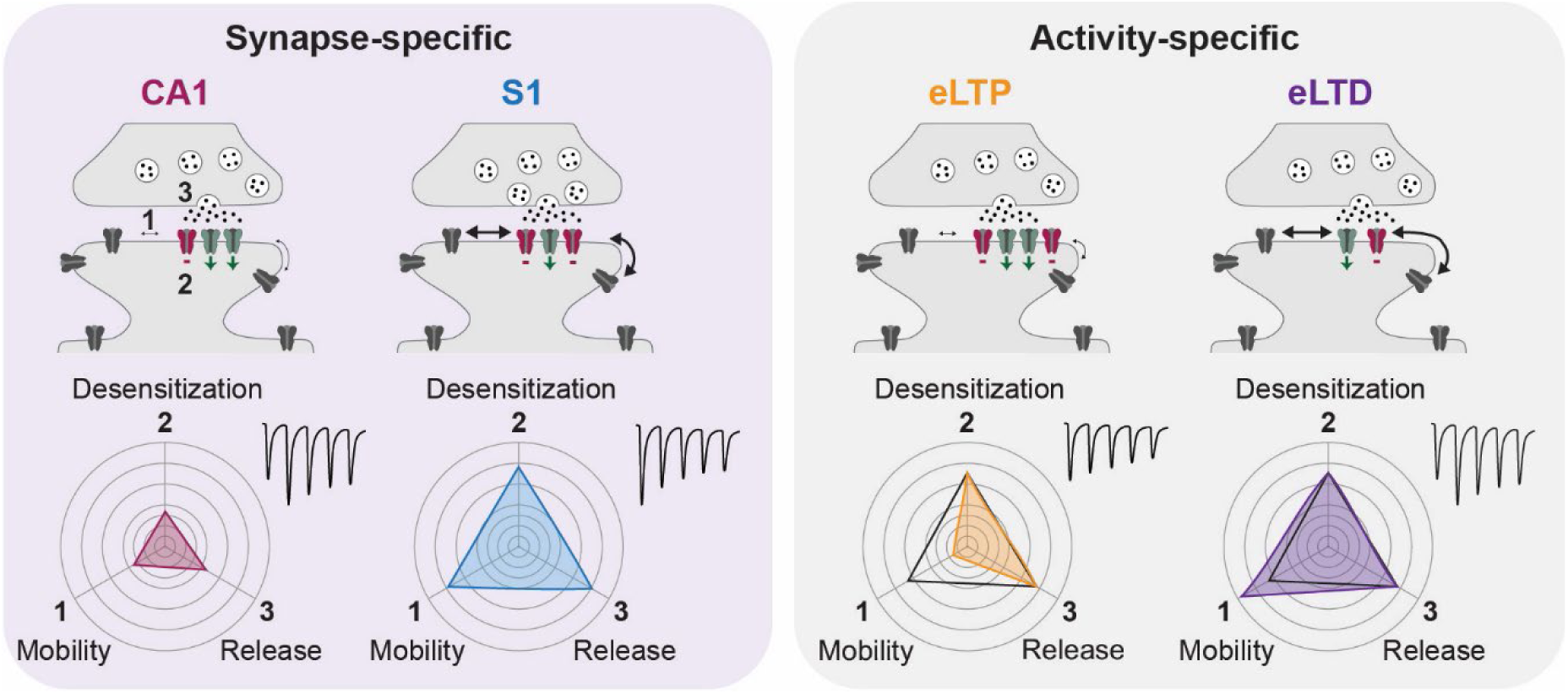
Model of the complementary roles of pre and post-synaptic parameters on HF-STP. The top row represents spine synapses in different conditions with 3 main parameters controlling STP. (1), AMPAR surface mobility – thicker arrows represent higher surface mobility. (2) AMPAR desensitization rate – red AMPAR are desensitized upon glutamate release. (3) Probability of glutamate release. The bottom row represents radar plots of these 3 parameters (from lower values in the inner circle to higher values in the outer circle). (1) Surface diffusion rate. (2) fraction of desensitized receptors after glutamate release (mixing rate of entry and exit from the desensitized state). (3) Probability of glutamate release. Left box represents the difference in contribution of these 3 parameters to STP in basal conditions in CA1 versus S1. Right box represents the difference between synapses having undergone eLTP or LTD.

## References

1 Zucker, R. S. & Regehr, W. G. Short-term synaptic plasticity. Annu Rev Physiol 64, 355–405, (2002).

2 Jackman, S. L. & Regehr, W. G. The Mechanisms and Functions of Synaptic Facilitation. Neuron 94, 447–464, (2017).

3 Jacobi, E. & von Engelhardt, J. Modulation of information processing by AMPA receptor auxiliary subunits. The Journal of physiology 599, 471–483, (2021).

4 Heine, M. et al. Surface mobility of postsynaptic AMPARs tunes synaptic transmission. Science 320, 201–205, (2008).

5 Frischknecht, R. et al. Brain extracellular matrix affects AMPA receptor lateral mobility and short-term synaptic plasticity. Nature neuroscience 12, 897–904, (2009).

6 Constals, A. et al. Glutamate-Induced AMPA Receptor Desensitization Increases Their Mobility and Modulates Short-Term Plasticity through Unbinding from Stargazin. Neuron 85, 787–803, (2015).

7 Opazo, P. et al. CaMKII triggers the diffusional trapping of surface AMPARs through phosphorylation of stargazin. Neuron 67, 239–252, (2010).

8 Compans, B. et al. NMDAR-dependent long-term depression is associated with increased short term plasticity through autophagy mediated loss of PSD-95. Nat Commun 12, 2849, (2021).

9 Christie, L. A. et al. AMPA receptor desensitization mutation results in severe developmental phenotypes and early postnatal lethality. Proc Natl Acad Sci U S A 107, 9412–9417, (2010).

10 Hansen, K. B. et al. Structure, Function, and Pharmacology of Glutamate Receptor Ion Channels. Pharmacol Rev 73, 298–487, (2021).

11 Zhang, D. et al. Structural mobility tunes signalling of the GluA1 AMPA glutamate receptor. Nature 621, 877–882, (2023).

12 Marvin, J. S. et al. Stability, affinity, and chromatic variants of the glutamate sensor iGluSnFR. Nat Methods 15, 936–939, (2018).

13 Getz, A. M. et al. High-resolution imaging and manipulation of endogenous AMPA receptor surface mobility during synaptic plasticity and learning. Science advances 8, eabm5298, (2022).

14 Rusakov, D. A., Savtchenko, L. P., Zheng, K. & Henley, J. M. Shaping the synaptic signal: molecular mobility inside and outside the cleft. Trends Neurosci 34, 359–369, (2011).

15 Groc, L. & Choquet, D. Linking glutamate receptor movements and synapse function. Science 368, (2020).

16 Nair, D. et al. Super-resolution imaging reveals that AMPA receptors inside synapses are dynamically organized in nanodomains regulated by PSD95. J Neurosci 33, 13204–13224, (2013).

17 Macgillavry, H. D., Song, Y., Raghavachari, S. & Blanpied, T. A. Nanoscale Scaffolding Domains within the Postsynaptic Density Concentrate Synaptic AMPA Receptors. Neuron 78, 615–622, (2013).

18 Tang, A. H. et al. A trans-synaptic nanocolumn aligns neurotransmitter release to receptors. Nature 536, 210–214, (2016).

19 Savtchenko, L. P. & Rusakov, D. A. Moderate AMPA receptor clustering on the nanoscale can efficiently potentiate synaptic current. Philos Trans R Soc Lond B Biol Sci 369, 20130167, (2014).

20 Bats, C., Groc, L. & Choquet, D. The interaction between Stargazin and PSD-95 regulates AMPA receptor surface trafficking. Neuron 53, 719–734, (2007).

21 Borgdorff, A. J. & Choquet, D. Regulation of AMPA receptor lateral movements. Nature 417, 649–653, (2002).

22 Tardin, C., Cognet, L., Batc, C., Lounis, B. & Choquet, D. Direct imaging of lateral movements of AMPA receptors inside synapses. The EMBO Journal 22, 4656–4665, (2003).

23 Polenghi, A. et al. Kainate Receptor Activation Shapes Short-Term Synaptic Plasticity by Controlling Receptor Lateral Mobility at Glutamatergic Synapses. Cell Rep 31, 107735, (2020).

24 de Luca, E. et al. Inter-Synaptic Lateral Diffusion of GABAA Receptors Shapes Inhibitory Synaptic Currents. Neuron 95, 63–69 e65, (2017).

25 Groc, L., Choquet, D. & Chaouloff, F. The stress hormone corticosterone conditions AMPAR surface trafficking and synaptic potentiation. Nat Neurosci 11, 868–870, (2008).

26 Makino, H. & Malinow, R. AMPA receptor incorporation into synapses during LTP: the role of lateral movement and exocytosis. Neuron 64, 381–390, (2009).

27 Penn, A. C. et al. Hippocampal LTP and contextual learning require surface diffusion of AMPA receptors. Nature 549, 384–388, (2017).

28 Chen, H., Roth, R. H., Lopez-Ortega, E., Tan, H. L. & Huganir, R. L. AMPA Receptors Exist in Tunable Mobile and Immobile Synaptic Fractions In Vivo. eNeuro 8, (2021).

29 Khodosevich, K. et al. Coexpressed auxiliary subunits exhibit distinct modulatory profiles on AMPA receptor function. Neuron 83, 601–615, (2014).

30 Petrini, E. M. et al. Endocytic trafficking and recycling maintain a pool of mobile surface AMPA receptors required for synaptic potentiation. Neuron 63, 92–105, (2009).

31 Schwenk, J. et al. Regional diversity and developmental dynamics of the AMPA-receptor proteome in the mammalian brain. Neuron 84, 41–54, (2014).

32 Shi, Y., Lu, W., Milstein, A. D. & Nicoll, R. A. The stoichiometry of AMPA receptors and TARPs varies by neuronal cell type. Neuron 62, 633–640, (2009).

33 Kamalova, A., Futai, K., Delpire, E. & Nakagawa, T. AMPA Receptor Auxiliary Subunit GSG1L Suppresses Short-Term Facilitation in Corticothalamic Synapses and Determines Seizure Susceptibility. Cell Rep 32, 107921, (2020).

34 Pampaloni, N. P., Riva, I., Carbone, A. L. & Plested, A. J. R. Slow AMPA receptors in hippocampal principal cells. Cell Rep 36, 109496, (2021).

35 Chamma, I., Levet, F., Sibarita, J. B., Sainlos, M. & Thoumine, O. Nanoscale organization of synaptic adhesion proteins revealed by single-molecule localization microscopy. Neurophotonics 3, 041810, (2016).

36 Carbone, A. L. & Plested, A. J. Coupled control of desensitization and gating by the ligand binding domain of glutamate receptors. Neuron 74, 845–857, (2012).

37 Sun, Y. et al. Mechanism of glutamate receptor desensitization. Nature 417, 245–253, (2002).

38 Reyes, A. et al. Target-cell-specific facilitation and depression in neocortical circuits. Nat Neurosci 1, 279–285, (1998).

39 Blackman, A. V., Abrahamsson, T., Costa, R. P., Lalanne, T. & Sjostrom, P. J. Target-cell-specific short-term plasticity in local circuits. Front Synaptic Neurosci 5, 11, (2013).

40 Rosenmund, C., Clements, J. D. & Westbrook, G. L. Nonuniform probability of glutamate release at a hippocampal synapse. Science 262, 754–757, (1993).

41 Hessler, N. A., Shirke, A. M. & Malinow, R. The probability of transmitter release at a mammalian central synapse. Nature 366, 569–572, (1993).

42 Silver, R. A., Lubke, J., Sakmann, B. & Feldmeyer, D. High-probability uniquantal transmission at excitatory synapses in barrel cortex. Science 302, 1981–1984, (2003).

43 Derdikman, D. et al. Layer-specific touch-dependent facilitation and depression in the somatosensory cortex during active whisking. J Neurosci 26, 9538–9547, (2006).

44 Campelo, T. et al. AMPAR-Dependent Synaptic Plasticity Initiates Cortical Remapping and Adaptive Behaviors during Sensory Experience. Cell Rep 32, 108097, (2020).

45 Götzke, H. et al. The ALFA-tag is a highly versatile tool for nanobody-based bioscience applications. Nat Commun 10, 4403, (2019).

46 Volianskis, A., Collingridge, G. L. & Jensen, M. S. The roles of STP and LTP in synaptic encoding. PeerJ 1, e3, (2013).

47 Shibata, A. C. E. et al. Photoactivatable CaMKII induces synaptic plasticity in single synapses. Nat Commun 12, 751, (2021).

48 Cook, S. G., Buonarati, O. R., Coultrap, S. J. & Bayer, K. U. CaMKII holoenzyme mechanisms that govern the LTP versus LTD decision. Sci Adv 7, (2021).

49 Abbott, L. F., Varela, J. A., Sen, K. & Nelson, S. B. Synaptic depression and cortical gain control. Science 275, 220–224, (1997).

50 Varela, J. A. et al. A quantitative description of short-term plasticity at excitatory synapses in layer 2/3 of rat primary visual cortex. J Neurosci 17, 7926–7940, (1997).

51 Hestrin, S. Activation and desensitization of glutamate-activated channels mediating fast excitatory synaptic currents in the visual cortex. Neuron 9, 991–999, (1992).

52 Wang, J. H. & Kelly, P. T. Regulation of synaptic facilitation by postsynaptic Ca2+/CaM pathways in hippocampal CA1 neurons. Journal of neurophysiology 76, 276–286, (1996).

53 Zhou, X., Tian, C., Zhang, X., Zheng, M. & Xu, K. Short-term plasticity as a mechanism to regulate and retain multistability. *Chaos*, Solitons & Fractals 165, 112891, (2022).

54 Wu, S. et al. Induction and propagation of transient synchronous activity in neural networks endowed with short-term plasticity. Cogn Neurodyn 15, 53–64, (2021).

55 Rao-Ruiz, P., Visser, E., Mitrić, M., Smit, A. B. & van den Oever, M. C. A Synaptic Framework for the Persistence of Memory Engrams. Front Synaptic Neurosci 13, 661476, (2021).

